# Biophysically realistic network-level transport model of tau progression with exosome-mediated release and uptake processes

**DOI:** 10.64898/2026.01.21.700898

**Authors:** Nuutti Barron, Veronica Tora, Emilia Cozzolino, Michiel Bertsch, Ashish Raj

## Abstract

The spatiotemporal progression of tau in neurodegenerative diseases like Alzheimer’s follows the brain’s structural connectome, yet a gap exists between the macroscopic spread observed over years and the protein kinetics occurring over hours. Current models fail to reconcile this disparity or incorporate the cellular mechanisms driving transmission. Here, we advance the Network Transport Model (NTM) to bridge these scales by integrating active transport along microtubules, continuous toxic tau production, and exosome-mediated release and uptake. This framework constitutes one of the most biologically detailed models of tau spread on a whole brain to date, representing a significant innovation in how multiscale proteinopathies are simulated.

Simulations on the mouse connectome demonstrate that this framework replicates empirical tau propagation patterns. Our results identify trans-neuronal release and uptake rates as the primary “bottleneck” on macroscopic spread, providing a biologically grounded explanation for the disease’s slow progression. Furthermore, we find that high aggregation sequesters tau within regions, limiting global transmission, while increasing the abundance of toxic polymeric tau fibrils. Meanwhile, tau transport polarity bias (anterograde vs. retrograde) dictates spatial patterning. By linking molecular mechanics to system-wide pathology, this model provides an “in-silico” framework to evaluate how cellular-targeted interventions might alter the trajectory of tauopathic dementias.

**Author Summary:** In Alzheimer’s disease and related dementias, a protein called tau forms misfolded aggregates and gradually spreads through the brain along its neuronal wiring. A long-standing puzzle is linking the microscopic protein dynamics taking minutes to hours to the multiyear macroscopic spread of the disease. We developed a new mathematical model that is capable to explain macroscopic spreading behavior of tau as an emerging process from the underlying biophysical mechanics of tau pathology. Our simulations reproduce real patterns of tau spread, and our framework will pave the way for testing how treatments targeting specific cellular mechanisms might reshape disease progression.

## 1 Introduction

Accumulation and deposition of abnormal microtubule-associated protein tau in the brain is a characteristic pathology of neurodegenerative tauopathies such as Alzheimer’s disease (AD) and Frontotemporal dementia (FTD). The spatiotemporal progression of tau presents in a consistently staged fashion, where neurofibrillary tangles, aggregates of toxic tau, appear in consistent spatial patterns at progressive time points throughout disease progression. In AD for instance, tau appears in abundance in the locus coeruleus, after which it appears in other areas of the brain, such as the hippocampus, entorhinal cortex, and temporal lobe, and later in the cortex [12, 11]. While there is significant heterogeneity in spatiotemporal patterns of tau progression between individuals and diseases, substantial evidence suggests that starting from an initial seed site, misfolded tau protein is transmitted between neurons across the synapse [56, 25] and can be shuttled via exosomes [53], perhaps mediated by glial cells such as oligodendrocytes [16] and microglia [9],. On the whole brain level these processes manifest as the spread of tau over time along neuronal white matter tracts interconnecting brain regions.

Uncovering the biophysical mechanism underlying disease progression, such as in AD and FTD, is crucial to develop treatments by revealing possible cellular mechanisms to target with therapeutics. Modeling based methods of tau spread mediated by the brain’s neuronal white matter tracts provide a framework to study empirically observed patterns of tau progression. These models typically employ a graph diffusion approach, where tau spread is approximated as a simple diffusive process that occurs on a graph or network representation of the brain called the structural connectome (SC) that represents gray-matter regions of the brain as nodes, and neuronal white matter tracts as edges weighted by neuronal connectivity strength. These models, such as the Network Diffusion Model (NDM) [37] and successive connectome based diffusion models [17, 24, 42, 55, 54, 32, 8, 4, 35] predict regional tau pathology at time points following an initial tau seed as a diffusive function of concentration gradients on the structural connectome network. SIR (Susceptible-Infected-Removed) and Epidemic Spread Models (ESM) models have similarly modeled tau progression on the structural connectome following the original proposal by Iturria-Medina [24], where tau is modeled as a population and transmission on the network as contagion spread. These models have subsequently been used successfully by other workers [57, 58, 44]. These models have shown remarkable success in capturing canonical Braak staging of AD [55, 54] [57] and empirically derived spatiotemporal presentation of tau in human subjects [38, 42, 39] and mouse models of disease [31, 4].

Despite these successes, diffusion based graph models of tau spread have several limitations that prevent them from becoming complete and biophysically realistic models. In general, they fail to capture the microscopic cellular mechanisms that mediate tau spread. First, diffusive spread is at odds with our prevailing understanding of tau transmission along axons via active microtubule-based transport [43, 49]. Second, there is an inherent contradiction between the extremely slow timescale (order of months to years) of macroscopic tau progression and the timescale of underlying protein kinetics, like aggregation and transport (order of hours to days) [43]. Third, the kinetics of protein aggregation are not typically incorporated in popular models of macroscopic spread of tau, with some exceptions [39].

A two-neuron multi-compartmental model of microscale tau dynamics was proposed by Torok, *et al*. to explain tau aggregation, fragmentation, diffusion, and transport dynamics involving microtubule-attached molecular motors in both anterograde (in the direction of axon polarity) and retrograde directions (against the direction of axon polarity). Recently a Network Transport Model (NTM) [46] extended these microscale dynamics to the full structural connectome network as a set of transport-reaction type partial differential equations (PDEs). Its mathematical properties have been analyzed in [6]. To reduce numerical complexity of coupled PDEs a quasi-static approximation was proposed [46], where the PDEs on each edge were approximated by their steady states, leveraging the slow vs fast time scale property. The NTM is therefore a suitable platform to achieve the missing link between microscale cellular dynamics mediating tau spread and the macroscale regional patterns of tau progression.

The purpose of this study is to advance the mathematical machinery necessary to obtain a more complete and biophysically realistic model of tau propagation on the structural connectome, by overcoming four key mechanistic deficiencies of the prior NTM:

1. **Trans-neuronal release and uptake**. Trans-synaptic transmission of tau across the neuronal cell membrane is a key process [56, 25] that was previously modeled using a barrier permeability parameter that attenuates the diffusivity of tau across two barrier segments along the axon - at the axon initial segment (AIS) and at the synapse. This approach was chosen due to its mathematical tractability, but was understood to be a phenomenological rather than mechanistically accurate process. Here we propose to overcome this limitation by introducing an explicit and biologically grounded mechanism [53] by which tau transits between intra- and extra-cellular spaces, driven by exosome-mediated processes of tau release (intra-to-extra-cellular) and uptake (extra-to-intra-cellular) [33].
2. **Continuous tau production**. The production of toxic tau is hypothesized to be linked to the hyperphosphorylation and separation of microtubule associated tau [18, 30]. The previous model assumption of tau seeding as a singular event is not supported by the empirical observation that the total tau burden in the brain increases over time. The proposed model takes this into account by allowing for a nonzero continuous production term in seed regions after the initial seeding.
3. **Mechanistic control of time scales**. Prior modeling attempts have sought to address the vastly different time scales of macroscopic and microscopic processes via an arbitrary scaling constant between them, and by introducing diffusion barriers at synapse and AIS. However, this is a phenomenological element unsupported by mechanistic studies. Here, we attempt to naturally recapitulate this timescale disparity by controlling the release and uptake processes, which are biologically grounded and amenable to experimental intervention.
4. **Separation of intracellular and extracellular environments**. Key neurodegenerative processes are thought to occur preferentially in intracellular or extracellular environments. Hyperphosphorylation-driven detachment of normal microtubule-associated tau occurs intracellularly [18, 30], while exosome mediated spread of tau and its internalization by microglia occur in the extracellular environment [1, 9]. Here, we provide a more robust framework to model such processes by compartmentalizing intracellular and extracellular environments.

The key aspects of the NTM are described in **Figure 1**. By accounting for these critical microscopic dynamics, the proposed new NTM model allows us to explore how the macroscale evolution of tau relates to microscale aggregation-fragmentation, diffusion-transport, uptake-release, and tau production processes in simulations of tau spread in the mouse brain. The full model involves large coupled PDEs and is computationally prohibitive. To aid computational tractability we provide a quasi-static approximation algorithm, based on the concept of two time scales suggested by Tora et al. [46].

**Figure 1.**
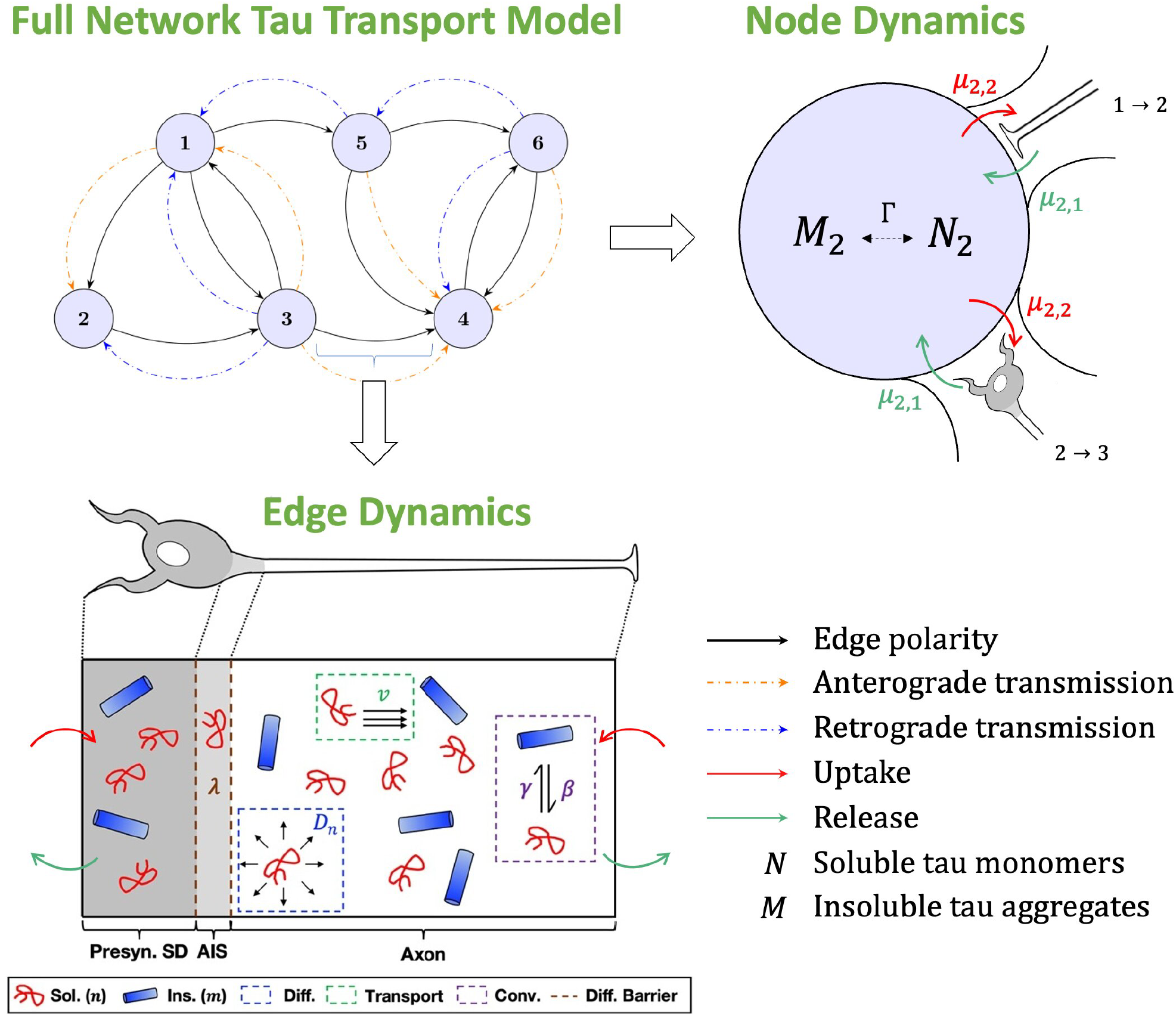
Illustration of the model system. At the whole network level, extracellular compartments are represented by nodes and white matter fiber projections between them by edges (top left panel). Tau pathology propagates on this network in an anterograde (orange dashed arrows) or retrograde (blue dashed arrows) direction, depending on the cell polarity and the properties of tau itself. Along the node-edge interface (top right panel, which portraits node 2 and its connections), tau undergoes the process of release (green arrows) from the intracellular to the extracellular space and uptake (red arrows), in the opposite direction. We use the active axonal transport model from Torok, *et al*. and cluster the synaptic and post-synaptic compartments in the endpoint node of the edge corresponding to the extracellular area where tau is released. The microscopic edge dynamics is schematized in the bottom panel. It models two distinct species of pathological tau, soluble (red) and insoluble (blue), within a multi-compartment system mimicking the single-edge system shown in the top panel. The main biological phenomena captured in this model are diffusion (blue box), active transport (green box), species interconversion through fragmentation and aggregation (purple box), and a diffusion-based barrier to inter-compartmental spread (brown dashed lines). Sol. - soluble tau; Ins. - insoluble tau; Diff. - diffusion; Conv. - tau interconversion; Diff. Barrier - diffusion barrier; Presyn. SD - presynaptic somatodendritic compartment; AIS - axon initial segment; SC - synaptic cleft; Postsyn. SD - postsynaptic somatodendritic compartment. Figure adapted from the original manuscript [49].

We performed model simulations on the whole-brain mouse connectome consisting of 426 regions, and numerical exploration of the behavior of its parameters. In contrast, prior models were limited to single neurons [49] or small subnetworks [46]. Although performing case-specific model fitting is currently computationally intractable using this framework, the NTM’s biophysical simulations demonstrate that the model replicates characteristic spatiotemporal patterns of tau propagation reported in empirical literature by comparing empirical end stage pathology to the spatial patterns of tau predicted by NTM simulations across a diverse set of biophysical NTM parameter regimes.

The new NTM model is capable of delivering more robust insights into how specific tau species may differentially propagate in the brain, governed by microscopic cellular mechanisms. We show that without active transport and protein kinetics these patterns could not emerge from earlier diffusion-based models. We also obtain mechanistically useful and sometimes surprising hypotheses about how each kinetic process governs eventual macroscopic progression pattern. We show, for example, that exosome release and uptake rates are the primary “brake” on the system, which presents a highly specific possible biological target to explore for drug intervention. Our simulations also strongly suggest that high tau aggregation rates may in fact prevent widespread transmission of tau by sequestering tangles within neurons. Finally, we explore model behavior in anterograde and retrograde transport regimes, which display strikingly different behavior in overall tau burden, rate of progression, staging and overall spatial patterning in simulations of tau progression in the mouse brain. Although this work focuses on understanding tau progression patterns in the mouse brain, it also establishes a framework to study tau progression in humans, and provides insight into the link between tau protein kinetics and global patterns of tau progression on a brain network that have possible clinical implications that can be useful for understanding and treatment of AD, FTD, and other tauopathic dementias. Thus our model provides a theoretical bridge that allows experimentalists to test how specific cellular interventions might alter the trajectory of disease across the entire brain.

## 2 Modeling

### 2.1 Overview

The brain is modeled as a network, referred to as the brain’s structural connectome, of a finite number of nodes and edges, where nodes represent brain compartments and edges the connections between them in terms of neuronal bundles. We present a Network Transport Model of how tau propagates on this network via active axonal transport. This model begins with the microscopic tau dynamics within a single neuron described by Torok, *et al*. [49]. Then it extends, similar to our earlier NTM model [46], tau spreading behavior across long-range white matter fiber projections. Refer to the illustration given in Figure 1. These dynamics provide a mathematical formulation for tau diffusion and directed active transport of soluble tau monomers, as well as aggregation and fragmentation of tau monomers and fibrils. This deviates from previous diffusion-based approaches, as it allows for directed spreading of tau aligned with or against neuronal polarity (from *pre-* to *post* -synaptic regions). These tau dynamics formed the basis of the edgewise tau spread behavior in the previously published NTM model [6], [46].

The present NTM model builds off of these edgewise dynamics, but fills four key mechanistic gaps (described in Introduction): a new exosome-mediated release and uptake process; continuous pathology production; mechanistic control of fast vs slow time scales; and separation of cellular compartments. The new model necessitated several noteworthy mathematical and implementation innovations. For instance, in the present version each edge is divided into three compartments instead of five, since we are no longer relying on phenomenological barriers in AIS and the synapse (see [49]). The exchange of soluble tau between nodes and edges is now modulated by newly introduced uptake and release mechanisms which require the introduction of suitable boundary conditions at the node-edge boundary.

The proposed model has the benefit of a much lower computational burden than the previous study [46]. This property follows from the hypothesis of slow release and uptake processes, which considerably simplifies the mass balance between nodes and their incident edges compared to the one arising in [46]. The resulting model is therefore more practical and versatile due to the reduction of complexity and computational costs. What follows is a formal definition of these dynamics in the newly proposed NTM. In Section 2.2 we introduce a single edge model which describes the dynamics of intracellular pathological tau, closely following the two-neuron axonal transport model by Torok *et al*. [49], the spreading of pathological tau along edges is modeled in terms of axonal transport dynamics in addition to passive diffusion. The main novelty of the current single edge model is the introduction of extracellular tau in the nodes and a description of the release and uptake process of pathological tau between the extracellular space in the nodes and the intracellular space on the edges. In Section 2.3 we define the whole brain graph, and in Section 2.4 we couple the single edge dynamics to the global network dynamics and obtain the Network Transport Model with release and uptake.

We take advantage of the experimentally observed separation of timescales in AD, i.e. a fast one for the kinetic and transmission processes and a slow one for the progression of tau pathology to obtain a meaningful acceleration in compute time, as in [46]. Thus it is natural to introduce a quasi-static approximation to the NTM, where the fast timescale becomes instantaneous with respect to the slow one (Section 2.5). Unlike the prior approach, here we were able to successfully relate the observed macroscopic separation of timescales to the specific microscopic mechanism of exosome-mediated release and uptake.

### 2.2 A single-edge model

We begin this section with a rather intuitive introduction to the most important aspects of the model and minimize the amount of mathematical formulas. From Section 2.2.1 we will be more precise.

We introduce some notation. A node will be indicated by *P*_*i*_ and the edge directed (according to its polarization) from *P*_*i*_ to *P*_*j*_ by *e*_*ij*_. We recall that an edge consists of bundles of neurons, but is represented by a one-dimensional single neuron (see Figure 1, bottom). For a more precise description of the graph we refer to Section 2.3.

The three key biological processes of tau pathology are: (1) aggregation-fragmentation of intracellular tau; (2) its spreading along brain connections by means of active transport in addition to passive diffusion; (3) the exosome-mediated exchange of soluble tau between the intracellular and the extracellular space, i.e. the release-uptake process which takes place at the node-edge boundaries. For the time being, all these processes will be described *in the fast timescale*.

Regarding point (2), active transport along axons is described by the (directionally-biased) velocity [49]

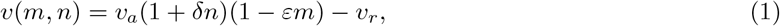

where *v*_*a*_, *v*_*r*_ *>* 0 are the baseline anterograde and retrograde velocities of tau, respectively; the parameters *δ, ϵ >* 0 govern the interactions between tau and the molecular motors (specifically kinesin) in the axon. In particular, the velocity (1) takes into account the fact that the propensity of soluble tau to travel in the anterograde direction is increased by a factor proportional to its concentration, *n*, and decreased by a factor proportional to insoluble tau concentration, *m*. Adding passive diffusion along the axon, this leads to the natural definition of a diffusion-transport flux of soluble tau from node *P*_*i*_ to *P*_*j*_ along edge *e*_*ij*_, indicated by *J*_*ij*_.

Concerning point (3), the exchange of soluble tau between intracellular and extracellular space is modeled by a suitable boundary condition for the fluxes *J*_*ij*_ at the vertex-edge boundaries. The flux along *e*_*ij*_ at node *P*_*i*_ is given by

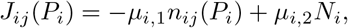

where *n*_*ij*_(*P*_*i*_) is the density of *intracellular* soluble tau along *e*_*ij*_ at the boundary point *P*_*i*_ of the edge, and *N*_*i*_ that of *extracellular* soluble tau at the node *P*_*i*_. Similarly, the flux along *e*_*ij*_ at the other node *P*_*j*_ is

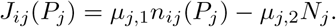

Here the parameters

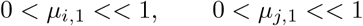

govern the release of soluble intracellular tau respectively at nodes *P*_*i*_ and *P*_*j*_; similarly, the parameters

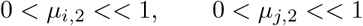

govern the uptake of soluble extracellular tau respectively at these nodes. In order to explain the existence of two very different timescales, we will assume that the release-uptake process strongly slows down the spreading of soluble tau along the network. Therefore, in the fast timescale the release-uptake parameters *µ*_*i,k*_ will be chosen relatively small compared to other parameters in the model. In Section 2.3, where the single-edge model is extended to the global network, we will see that *µ*_*i,k*_ are, in a first approximation, of the same order as the proportion of the observed timescales, i.e. the separation of timescales is determined by the microscopic properties of the release and uptake mechanism.

In the following sections we shall make the single-edge model, of which we discussed only some aspects in the present section, mathematically precise.

#### 2.2.1 The PDE-model

We describe the edge *e*_*ij*_ from *P*_*i*_ to *P*_*j*_ as a single neuron and introduce a 1D variable *x* ∈ [0, *L*_*ij*_]. Here *L*_*ij*_ is the total “length” of the neuron, which, for simplicity, is assumed to be constant for all *e*_*ij*_. Biological compartments within the neuron with distinct tau dynamics are represented by three distinguished segments (see Figure 1, bottom):

i. **Presyn. SD**: presynaptic somatodendritic compartment, (0, *x*_1_).
ii. **AIS**: axon initial segment, (*x*_1_, *x*_2_).
iii. **Axon**: axonal component, (*x*_2_, *L*_*ij*_).

Let *t*_fast_ denote time in the fast scale. Let *m*_*ij*_(*x, t*_fast_) and *n*_*ij*_(*x, t*_fast_) denote the densities at time *t*_fast_ per unit volume of insoluble and soluble intracellular pathological tau, respectively, on the edge *e*_*ij*_. Furthermore, *N*_*i*_(*t*_fast_) and *M*_*i*_(*t*_fast_) denote, respectively, the density of extracellular soluble and insoluble tau at vertex *P*_*i*_. The governing equation of *m*_*ij*_(*x, t*_fast_), *n*_*ij*_(*x, t*_fast_) on the edge *e*_*ij*_ are given by

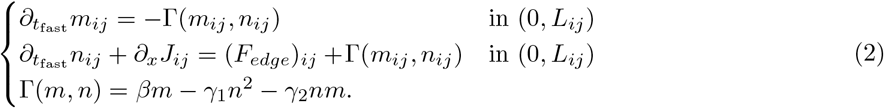

Here, ±Γ(*m, n*) are reaction terms taking into account interconversion between soluble and insoluble tau species, where *β* is the unimolecular rate of fragmentation, *γ*_1_ is the bimolecular rate of soluble-soluble tau aggregation, *γ*_2_ is the bimolecular rate of soluble-insoluble tau aggregation; *F*_*edge*_ denotes a source term for soluble pathological tau on the edge to model tau-seeding and is of the form

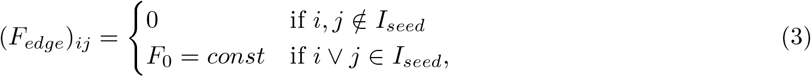

where *I*_*seed*_ is a set of indices corresponding to brain regions where tau pathology is assumed to initiate; the flux *J*_*ij*_ of *n*_*ij*_ on *e*_*ij*_ is, for fixed time, given by

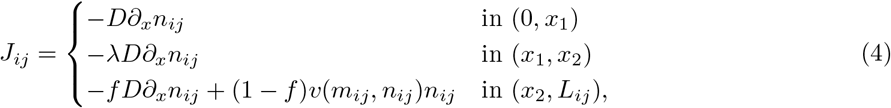

where, as in [49], *f* is the average fraction of soluble pathological tau along axons that is undergoing diffusion as opposed to active transport at any given time [27, 15], *λ* ∈ (0, 1) is the reduction of diffusivity in the AIS. Because *m*_*ij*_ represents the concentration of insoluble aggregates of tau, the diffusive and active transport flux of this species vanishes along edges.

As explained in the previous section, the exchange of soluble tau between intracellular and extracellular space is modeled introducing a suitable boundary condition at the vertex-edge boundary. In particular, system (2) is coupled with the following Neumann boundary conditions at *x* = 0 and *x* = *L*_*ij*_:

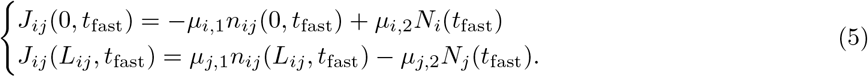

We stress that all coefficients introduced above refer to the fast timescale.

#### 2.2.2 Quasi-static approximation on edge

Let *ϕ <<* 1 be the proportion of the observed slow and fast timescales and set

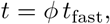

i.e. *t* denotes time in the slow timescale. We rewrite system (2)-(5) in the slow timescale. By abuse of notation, we use the same letters *n*_*ij*_, *m*_*ij*_, *N*_*i*_, *M*_*i*_ for the densities expressed in *t* instead of *t*_fast_. The same holds for the flux *J*_*ij*_ in the fast timescale, given by (4), i.e. *J*_*ij*_(*x, t*) expresses the flux in the fast timescale expressed in the slow timescale *t*. Then we obtain the system

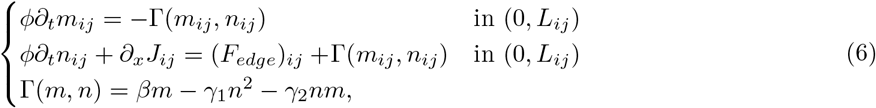

where *J*_*ij*_ is still given by (4), coupled to

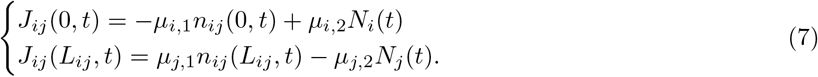

Since *ϕ* is a very small number, we approximate system (6) by setting *ϕ* = 0. This yields the quasi-static approximation of system (6):

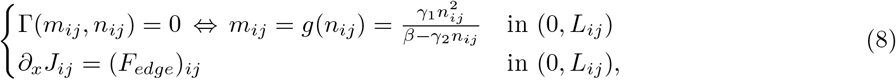

where *J*_*ij*_ is given by (4). Again (8) must be coupled to (7).

Formally, the quasi-static approximation is justified by the fact that, for each fixed time *t*, the equilibria of (6)-(7) are stable and, in the slow timescale, will be reached almost instantaneously due to the extreme smallness of *ϕ*.

Before extending the single-edge problem to the global network, we specify what we mean by the weighted connectivity graph.

### 2.3 The structural connectivity graph

The white matter tract connective wiring of the brain, on which tau is theorized to spread, can be described as a structural connectivity graph *G. G* is defined such that graph edges represent white matter tracts and nodes represent brain regions under a specified brain atlas parcellation. More precisely, *G* is defined as a weighted, directed graph with a finite number of vertices *P*_*i*_ and edges *e*_*ij*_ (*i* ?= *j*) directed from vertex *P*_*i*_ to *P*_*j*_. Edges *e*_*ij*_ and *e*_*ji*_ are defined by their polarization (or empirically obtained directionality): the polarization, or directionality, of *e*_*ij*_ goes from *P*_*i*_ to *P*_*j*_, whereas *e*_*ji*_ is directed from *P*_*j*_ to *P*_*i*_.

The graph edge weights are defined by a weight function *c* such that:

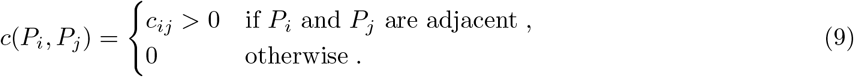

The connectivity weights *c*_*ij*_ express the empirically obtained white matter connectivity strength between the *i*-th and *j*-th discrete brain regions. Note that in a directed graph the weight function is not required to be symmetric; the weight *c*_*ij*_ can be different from *c*_*ji*_. The connectivity strength is an experimentally obtained measure proportional to the number of axons within a white matter tract, which is related to its cross-sectional area. This definition of connectivity strength has been widely used within diffusion-based mathematical models developed by our group [37], [4], [8] and others [24], [55], [42]. A critical distinction here is that graph nodes are defined to represent the extracellular space of the associated brain regions, and that graph edges represent the intracellular space of white matter tract neurons connecting brain regions. Thus, cellular uptake and release processes represent the exchange of tau between the extracellular space of a brain region and the intracellular space of the white matter tracts synapsing in the brain region. Here we utilize the mesoscale mouse connectivity atlas (MCA) from the Allen Institute for Brain Science [34], which uses viral tracing to determine both the connectivity strength (connectome edge weights) and polarity (edge direction) of mouse brain white matter tract connections (connectome edges) between region-pairs at a fine regional parcellation (utilizing 426 brain regions). The directness of such connections has been well-established. See [34] for more details. We simulate the NTM on the entire mouse brain network, building on prior studies that used the NTM and focused only on a brain sub-circuit consisting of 11 hippocampal structures, 3 retrosplenial areas from the neocortex, and the piriform area (for a total of only 30 regions across both hemispheres) [46].

Concerning the choice of the weight *c*_*ij*_, we refer to [46]; it measures the connectivity densities of white matter fiber projections obtained from the mouse structural connectome (see (9)). We remind that this choice was made in order to represent mass flow along the edge *e*_*i,j*_ by the term *c*_*ij*_*J*_*ij*_. Such modeling assumption is supported by tractography literature [34], but it is beyond the current scope to establish an exact equivalence between the chosen measure of connection strength (i.e. tracer-based) and axonal density.

### 2.4 The global Network Transport Model (NTM) with release and uptake

We begin with the PDE-model. Let *M*_*i*_(*t*) and *N*_*i*_(*t*) denote the densities at time *t* per unit volume at vertex *P*_*i*_ of, respectively, insoluble and soluble extracellular pathological tau protein, where *t* refers to the slow time scale. The governing equations for *M*_*i*_ and *N*_*i*_ are given by

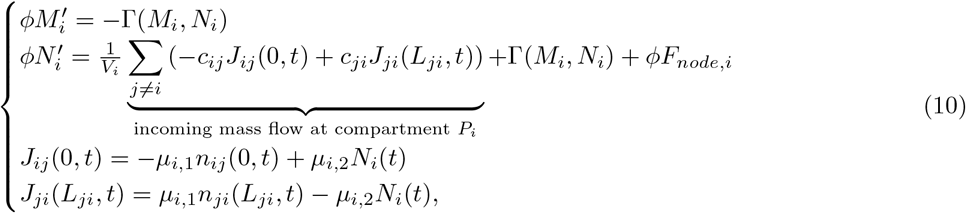

coupled to system (6)-(7) for *m*_*ij*_ and *n*_*ij*_. Here *ϕ <<* 1 is the proportion of the two timescales, 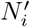 and 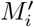 denote derivatives with respect to *t*, the flux *J*_*ij*_(*x, t*) is given by (4), Γ(*M, N*) is defined as in (6) (with coefficients *β, γ*_1_ and *γ*_2_ referring to the fast timescale), *V*_*i*_ is the volume of the brain compartment *P*_*i*_, and the production term *F*_*node,i*_ describes, in the slow timescale, the recruitment and accumulation of extracellular tau pathology in the brain regions where tau pathology initiates:

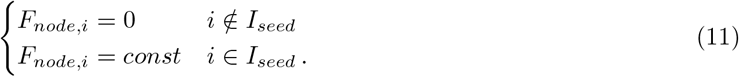

The incoming flux of soluble tau into node *P*_*i*_ with respect to the fast timescale,

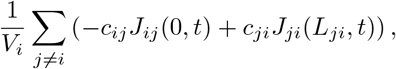

describes the mass transfer mechanism of soluble tau between the node *P*_*i*_ and the edges *e*_*ij*_ and *e*_*ji*_. Combined with the expressions for *J*_*ij*_(0, *t*) and *J*_*ij*_(*L*_*ij*_, *t*) it models the release and uptake mechanism, i.e. the exchange of pathological tau between the intracellular and extracellular space.

The PDE-model with release and uptake is completed by initial data for *N*_*i*_(0) and *M*_*i*_(0).

### 2.5 Quasi-static approximation

Since *ϕ <<* 1, we proceed as in Section 2.1.2 to approximate the PDE-model by its quasi-static approximation, where the fast timescale becomes instantaneous in the slow one. The first equation in (10) leads, in the quasi-static approximation, to

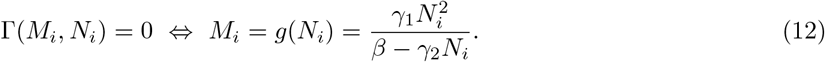

In addition, given *N*_*i*_(*t*) in the nodes *P*_*i*_, *n*_*ij*_(*x, t*) and *m*_*ij*_(*x, t*) of (2) must satisfy (7)-(8) in the quasi-static approximation, as discussed in Section 2.1.2.

In view of (12), in the slow time scale the quasi-static mass balance only needs to take into account the production term and the fluxes between the brain compartment *P*_*i*_ and the edges *e*_*ji*_ and *e*_*ij*_. By (7) this leads to

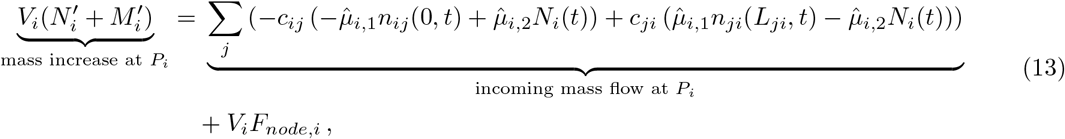

where we have set

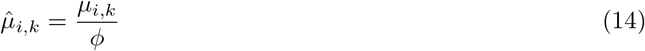

to take account that the incoming mass flow must be taken in the slow timescale. Finally, we substitute *M*_*i*_ = *g*(*N*_*i*_) in (13) to calculate 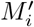 and obtain the equation for *N*_*i*_:

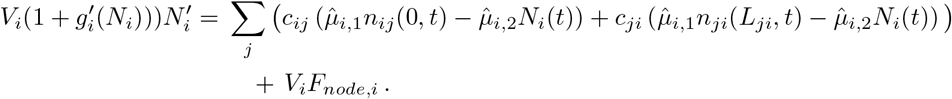

#### Remark 1

*Observe that, in* (14), *ϕ is the observed proportion of timescales and the coefficients µ*_*i,k*_ *reflect the microscopic properties of the release and uptake mechanism. By definition of the slow timescale we require that* 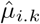 *is a coefficient of order 1, i*.*e. that µ*_*i,k*_ *is of order ϕ. This means that our modeling of the release and uptake mechanism suggests a relation between the order of magnitude of the slow timescale and the coefficients µ*_*i,k*_.

Reassuming, the governing equations of the quasi-static model for pathological tau are given by

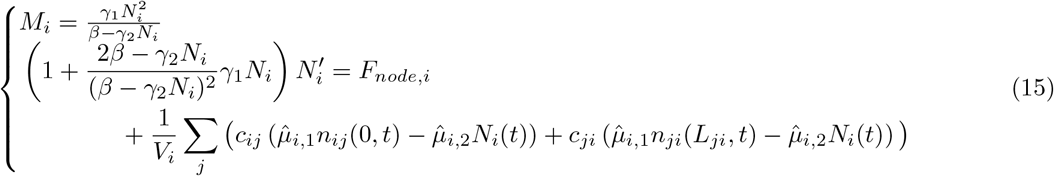

coupled to (7)-(8) and given initial data *N*_*i*_(0) = *N*_0*i*_ at each mode *P*_*i*_.

As in [46], we observe that, if *γ*_2_ *>* 0, the function *g* in (12) has a singularity at *n* = *β/γ*_2_. Under some natural constraints on the initial total mass of the system, it is possible to show that, given 0 ≤*N*_0*i*_ *< β/γ*_2_, the quasi-static model with release and uptake possesses a unique subcritical solution.

The proof of this statement will be presented in a separate paper.

### 2.6 Model implementation

#### 2.6.1 Implementation of the quasi-static edge problem

In order to implement the quasi-static NTM with release and uptake (15), we first need to calculate for any edge the steady state solution *n*_*ij*_ evaluated at the points *x* = 0, *x* = *L*. Integrating the quasi-static edge equation (8)_2_ and keeping in mind equation (4) we see that

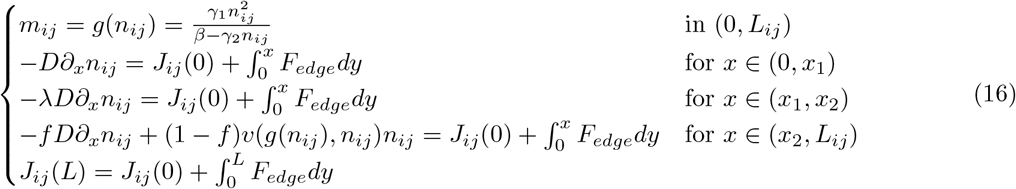

with boundary conditions

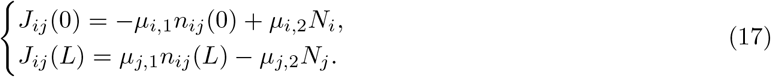

Setting *B* := *n*_*ij*_(0), we can define *n*_*ij,B*_(*x*) as the steady state profile expressed as a function of its initial data; then for given *N*_*i*_, *N*_*j*_ we solve the following problem: find *B* ∈ ℝ ^+^ such that

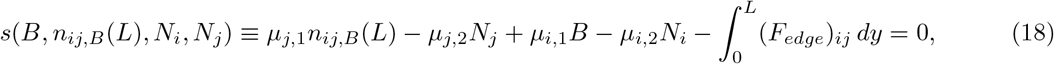

where *n*_*ij,B*_ satisfies the Cauchy problem

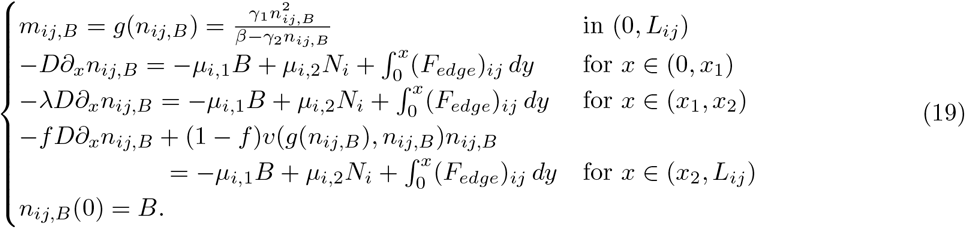

For given *N*_*i*_, *N*_*j*_, problem (19) on the edge *e*_*ij*_ is first integrated on an inhomogeneous spatial grid as a function of the shooting parameter *B* through the MATLAB function ode45, a solver based on an explicit Runge-Kutta method of order (4,5); then the shooting problem (18) is solved using the MATLAB’s nonlinear solver fsolve. Once the shooting parameter *B* := *n*_*ij*_(0) is calculated, the steady-state solution *n*_*ij*_(*x*) is also determined to obtain its value at *x* = *L*.

#### 2.6.2 Implementation of the quasi-static NTM with release and uptake

The quasi-static approximation to the NTM with release and uptake (15) is implemented in MATLAB version 2024b following Algorithm 1.

##### Algorithm 1

Quasi-static *Release-Uptake-NTM* Numerical Scheme

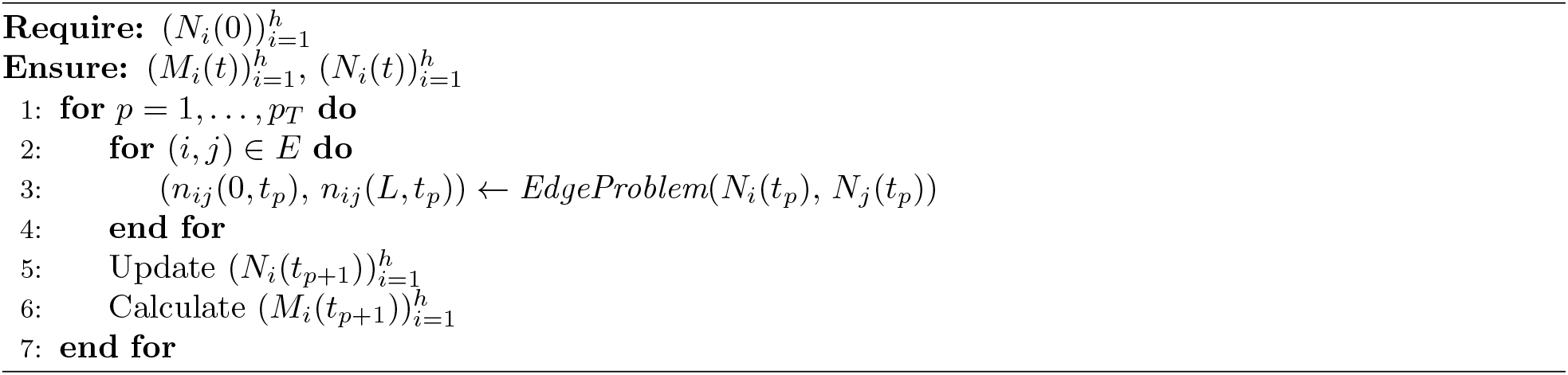

Introducing a inhomogeneous time grid of size *p*_*T*_ : *t*_*p*_ = *p*Δ*t, p* = 0, 1, 2, …, *p*_*T*_, given the node concentrations 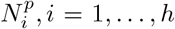 at time step *p*, the quasi-static edge problem (19) is solved on each edge *e*_*ij*_ ∈ *E* as described in section 2.6.1. This procedure is computed by the *EdgeProblem* function, which produces as output the edges concentrations *n*_*ij*_(0, *t*_*p*_), *n*_*ij*_(*L, t*_*p*_) at time *t*_*p*_ for all edges *e*_*ij*_ ∈ *E*. Then, the node concentration 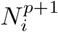 at time *t*_*p*+1_ is calculated at each node *P*_*i*_ ∈ *V* as a function of the edges concentrations *n*_*ij*_(0, *t*_*p*_), *n*_*ji*_(*L*_*ji*_, *t*_*p*_) at time *t*_*p*_ according to a first-order Euler discretization scheme of (15). To increase accuracy of the discretization scheme along the initial stages of evolution, we perform an intermediate mid-point approximation of (15) between successive time steps.

We then express the edge problem (16)-(17) as function of its parameters *λ, γ*_1_, *ϵ, δ, F*_*edge*_, *µ*_1_, *µ*_2_ expressed in their natural timescale (the fast one); more precisely, for any edge *e*_*ij*_ we define the steady state profiles as *n*_*ij*_ := *n*_*ij*_(*x, λ, γ*_1_, *ϵ, δ, F*_*edge*_, *µ*_1_, *µ*_2_). Analogously, the node concentrations can be defined as 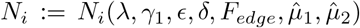, where also the edge parameters are included due to the dependence of the *N*_*i*_ on the profiles *n*_*ij*_ evaluated at *x* = 0, *x* = *L*. Then, fixing a suitable range for the parameters 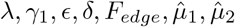, the above described algorithm is run to obtain the nodes concentration *N*_*i*_ along the entire time grid for any given parameter set. Observe that by (15)_1_ the concentration of insoluble extracellular tau (*M*_*i*_) is obtained once the computation of *N*_*i*_ is complete. The simulations are run in parallel using the computational resources provided by the University of California, San Francisco.

In order to provide a global picture of the evolution of tau pathology on the whole brain connectome network, we first define the total tau burden at each the vertex of the network as:

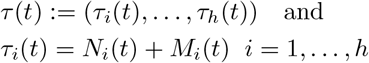

where *h* is the number of vertices of the graph *G* (see **Section 2.1**). However, we differentially investigate the nodal values *N*_*i*_ and *M*_*i*_ as each play a different role in the process of spread and neurodegeneration related to tau. We chose the Field CA1 in the left hemisphere of the mouse brain as our initiation site for tau pathology, as it matches the tau seed injection site for a number of empirical datasets of tau propagation from mouse studies we can compare NTM model simulations to. Mathematically, this can be expressed in the following way:

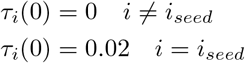

where *i*_*seed*_ is the index corresponding to the Field CA1 Left Hemisphere seed site. Without loss of generality, we set the initial tau burden to be 0.02 at the seeding location. Although we do not perform data fitting at this stage, our theoretical model focuses on P301S mouse models. Indeed, the mice data we compare NTM-predicted end-stage pathology for this type of tau. Observed significant variability in spread in these P301S models motivates the use of mathematical models to explain these differences. To explore the model dynamics, we ran a grid search on six key model parameters: 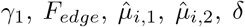, and *ϵ*. It has been previously shown that the ratio of aggregation rate to fragmentation rate governs the distribution of tau for the single-edge model [49], and therefore we hold the fragmentation rate, *β*, constant. The diffusivity barrier parameter *λ* showed to have little effect on NTM simulations, so this was fixed in numeric simulations presented here as well. To accurately model biophysics of spontaneous toxic tau production, we explore the *F*_*edge*_ parameter and set *F*_*node*_ = 0 everywhere, as tau production is hypothesized to occur in the intra-cellular environment captured by on connectome edges (instead of the extracellular environment captured by connectome nodes). As mentioned above in **Section 2.5**, the system exhibits a singularity for *γ*_2_ *>* 0, therefore we set *γ*_2_ = 0 for all simulations. For simplicity, we also require the release and uptake coefficients to be constant on the network, i.e. 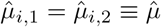 for all nodes *P*_*i*_.

### 2.7 Empirical Datasets

We compare NTM simulations of tau progression to 12 empirical mouse datasets of tau spread. Each selected empirical dataset abided by the following criteria: (1) regional tau abundance was quantified or semi-quantified based on immunochemistry; (2) the P301S tau mutant mouse model was used to prevent conflating endogenous tau protein types with other analyses; (3) had a minimal spatial coverage of a minimum of 40 brain regions with tau abundance quantified; (4) tau abundance was recorded at a minimum of three different time points. These same empirical tau datasets in mice have been used in studies of connectome-mediated spread of tau in previous work [4, 49, 50, 47]. The 12 mouse tauopathy datasets are referred to here as *IbaP301S* [22], *IbaHippInj* [21], *IbaStrInj* [21], *Hurtado* [20], *DS4* [26], *DS6* [26], *DS7* [26], *DS9* [26], *DS6 110* [26], *DS9 110* [26], *BoludaDSAD* [10], and *BoludaCBD* [10], and originate from five studies of tau progression in P301S mouse models. For more details, see Supplementary Table S1, previous connectome-mediated tau spread modeling work in mice using these datasets [4, 49, 50, 47] or the original studies [21, 22, 20, 26, 10].

## 3 Numerical simulations

Each NTM simulation uses the 426 region mesoscale connectivity atlas (MCA) of the mouse brain obtained from Oh et al. [34] for a high resolution abstraction of the brain’s graph network. However, for visualization purposes, we use a more coarse 44-region parcellation of the mouse brain used as the native space for sampling empirical tau data in Kaufman, et al. [26] referred to here as the ‘DS’ atlas when plotting tau trajectories (regional concentration over time). Glass brains utilize the finer-grade MCA atlas used in NTM simulations. To convert tau simulations from the 426-region MCA to the 44-region DS atlas for visualization, we compute the volume corrected mean tau concentration of all MCA atlas regions contained in a single DS atlas region at each simulated time point.

We examine the dynamics of the NTM under four different biophysical regimes: 1) under low and high tau aggregation conditions by varying the kinetic aggregation rate (governed by *γ*); 2) under anterograde and retrograde biased active tau transport conditions (governed by *δ* and *ϵ*); 3) under a range of tau uptake and release rates between white matter neuron’s intracellular and brain regions’ extracellular environments (modulated by 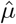); and 4) under low, intermediate, and high intracellular seed tau production conditions (modulated by *F*_*edge*_). For each comparison, the parameter modulating the biophysical mechanism of interest is varied while the remaining NTM parameters are held constant in order to examine the macroscale impact of modulating the microscale mechanisms governing tau dynamics in the NTM. We specifically focus on a few key characteristics of spatiotemporal tau dynamics: 1) the global rate of tau spread on the brain connectome network, 2) the staging of regional tau pathology, where the ordering of regions based on peak tau concentrations and their arrival times (how long it takes for a region to reach its peak concentration), and 3) the observed tau burden over time, described as the total amount of tau protein in the brain over time. We primarily examine the progression of soluble tau (*N*_*i*_ as described in the modeling section above) because the quasi-static NTM approximation computes insoluble tau concentration (*M*_*i*_ as described in the modeling section) as a monotonic function of the soluble tau concentration scaled by the fragmentation rate parameter *γ*_1_ when the *β* fragmentation rate parameter is fixed and *γ*_2_ = 0 as in the numeric simulations we explore (Equation 12). This means that the effects of NTM parameters on spatiotemporal patterns of soluble tau will carry over to insoluble tau, with the exception of *γ*_1_. The effect of *γ*_1_, referred to henceforth simply as *γ*, on the balance between soluble and insoluble tau is explored below.

### 3.1 Varying tau aggregation rate

Figure 2 demonstrates the effect of modulating the microscopic tau aggregation rate on macroscopic, whole-brain level spatiotemporal evolution of tau. Under low aggregation conditions, where *γ* = 0.001, tau concentration over time plots and heatmaps (Figure 2A, C; left) display a rise to a peak concentration value between 1 and 4 months followed by a decrease until the concentration plateaus to an equilibrium state with a homogeneous growth rate across regions. The seed region is unique in that it displays a decrease from the initial seed concentration to the end state with no peak. Notably, the time point at which the peak concentration for non-seed regions occurs varies between regions. For example, the concentration of tau in Field CA3 in both the left and right hemisphere peaks at around 2 months, while the concentration in the Dentate Gyrus and Field CA1 in the right hemisphere peak after 3 months. In contrast to low aggregation rate conditions that exhibit transient tau concentration peaks in non-seed regions, regional tau trajectories in non-seed regions generally exhibit a monotonic increase over time under high aggregation conditions (Figure 2A, C; right). This points to the fact that tau aggregation rate influences tau staging. The fact that high aggregation conditions do not demonstrate peak concentrations within the 12 month span of the simulation also points to increased aggregation slowing the spread rate of tau. The intuition behind this is that soluble tau acts as the conduit by which tau spreads on the connectome network, and high tau aggregation means a greater proportion of tau will convert to the insoluble filament form, which does not exhibit spread dynamics in the NTM. In other words, higher aggregation sequesters a greater proportion of the total pathologcial tau that is able to spread on the network, and with less tau available for spreading, the slower the spread on the network will be.

In addition, the lower aggregation rate simulation displays increased soluble tau burden (Figure 2B, left), as the concentration levels of tau across all regions is greater than in the high aggregation rate condition. This may appear to be a counterintuitive result, as generally it is thought that increasing tau aggregation increases the amount of toxic tau in the brain. However, this is the expected behavior for *soluble* tau on the network, as decreasing the tau aggregation rate decreases the amount of oligomeric soluble tau that is converted to polymeric insoluble tau fibrils, increasing the amount of tau that exists in soluble form. The opposite effect is observed when examining the concentration of polymeric *insoluble* tau filaments between high and low aggregation conditions as shown in Figure 3. Under low aggregation conditions (Figure 3A) the insoluble tau concentration in the brain is notably lower than under high aggregation conditions (Figure 3B). This distinction matches the current understanding that insoluble tau filaments are strongly associated with neurodegeneration in tauopathies such as AD and that increased tau aggregation increases the burden of these filaments. Overall, we conclude that aggregation dynamics affect patterns of both tau staging and overall spread rate, but have a much stronger impact on modulating global tau burden of both soluble and insoluble tau in the brain.

**Figure 2.**
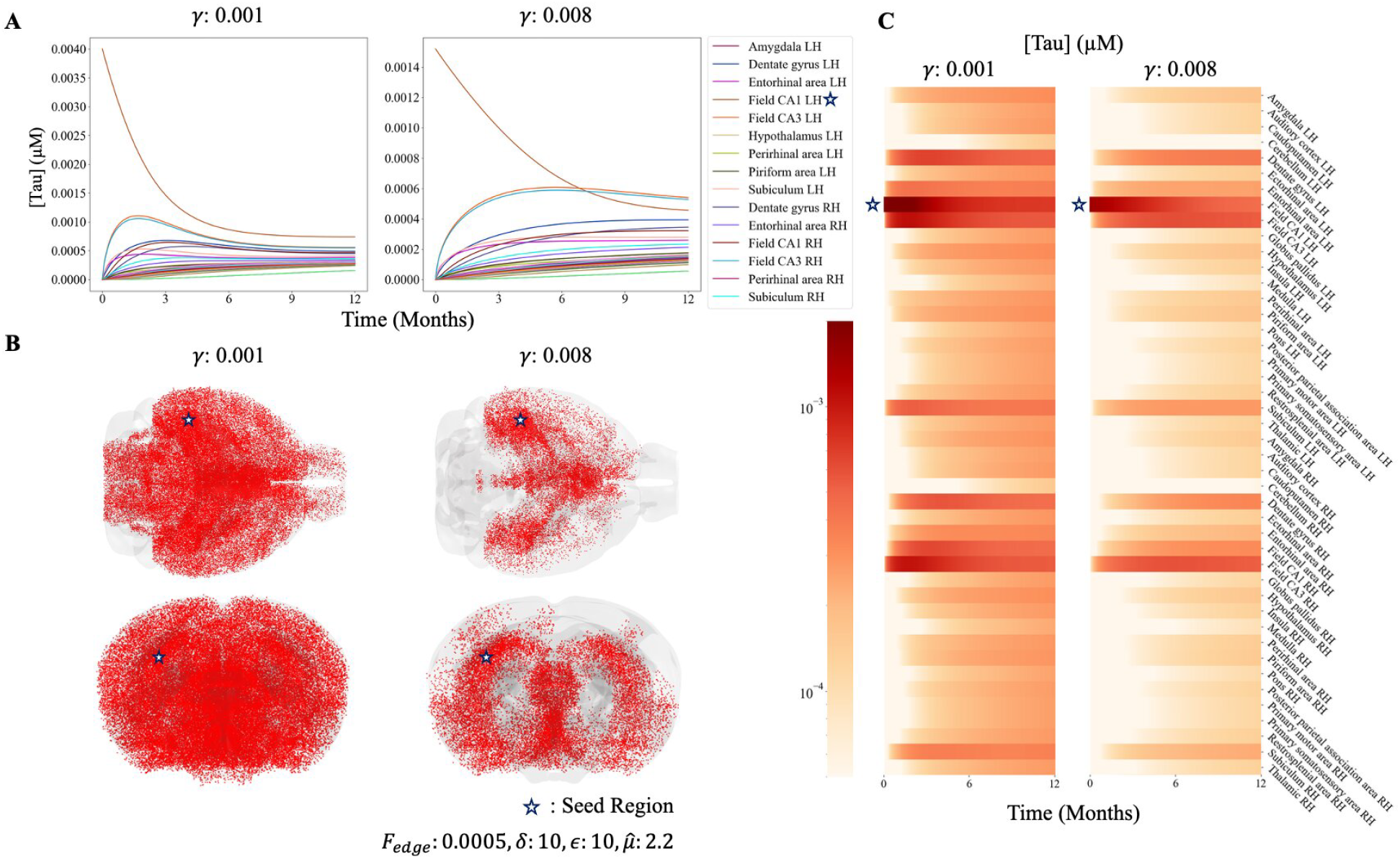
Effect of tau aggregation on NTM simulations. **A.** Soluble tau concentration over time plots for a 44 region parcellation of the mouse brain (converted from the 426 region MCA atlas used in NTM simulations) under low tau aggregation (left, *γ* = 0.001) and high aggregation (right, *γ* = 0.008) conditions. **B**. Glass mouse brains showing spatial distribution of soluble tau in the brain at 6 months post seeding with low aggregation (left) and high aggregation (right) conditions. Axial (top) and coronal (bottom) angles are shown, where a higher density of red dots indicates a higher tau concentration. **C**. Heatmap representation of per-region soluble tau concentration over time shown in **A**. The seed region (Field CA1, Left Hemisphere) is denoted with a blue star.

**Figure 3.**
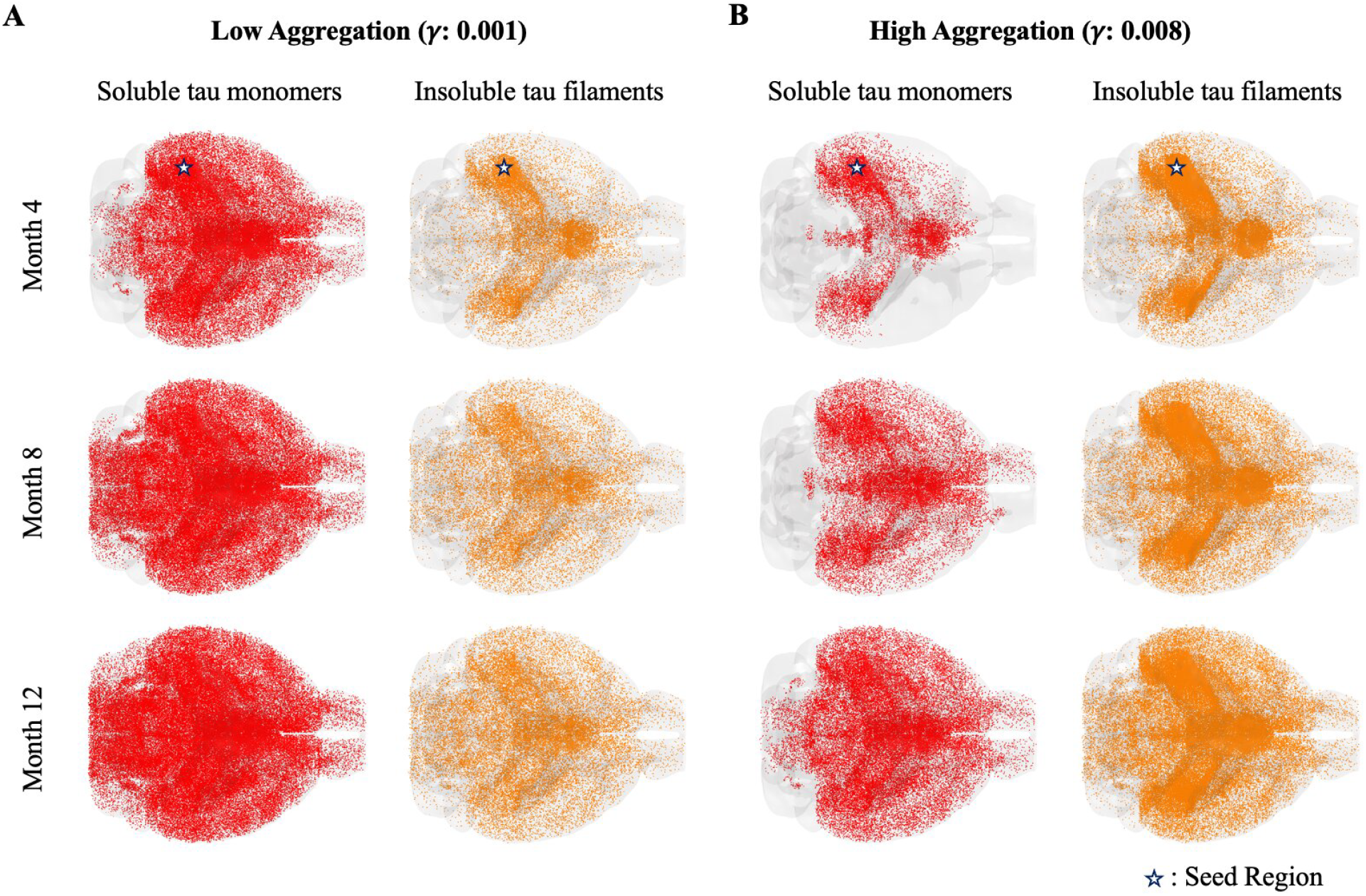
Spatial patterns of soluble versus insoluble tau under different aggregation conditions. Glass mouse brains displaying spatial distribution of tau under (**A**) low aggregation and (**B**) high aggregation conditions at four month intervals following the initial tau seeding. For each aggregation condition both soluble monomeric tau (left, red) and insoluble polymeric tau fibril (right, orange) concentrations are displayed. An greater density of dots indicates a greater concentration. NTM parameters and seeding is the same as Figure 2. The seed region is denoted with a blue star in the top row of glass brains.

### 3.2 Varying molecular motor rates

The *δ* and *ϵ* parameters of the NTM are designed to control the rate of kinesin transport of tau in the intracellular environment along axonal microtubules, where *δ* increases kinesin transport as a function of *soluble* oligomeric tau concentration and where *ϵ* decreases kinesin transport as a function of *insoluble* polymer tau concentration. Thus an anterograde or retrograde bias in the directional active transport of tau within neurons can be introduced by modulating *δ* and *ϵ* [49, 46]. *This does not have a notable impact on overall tau burden (Figure 4C; top two rows), which is expected given that modulating transport dynamics should not increase or decrease global levels of soluble tau*.

**Figure 4.**
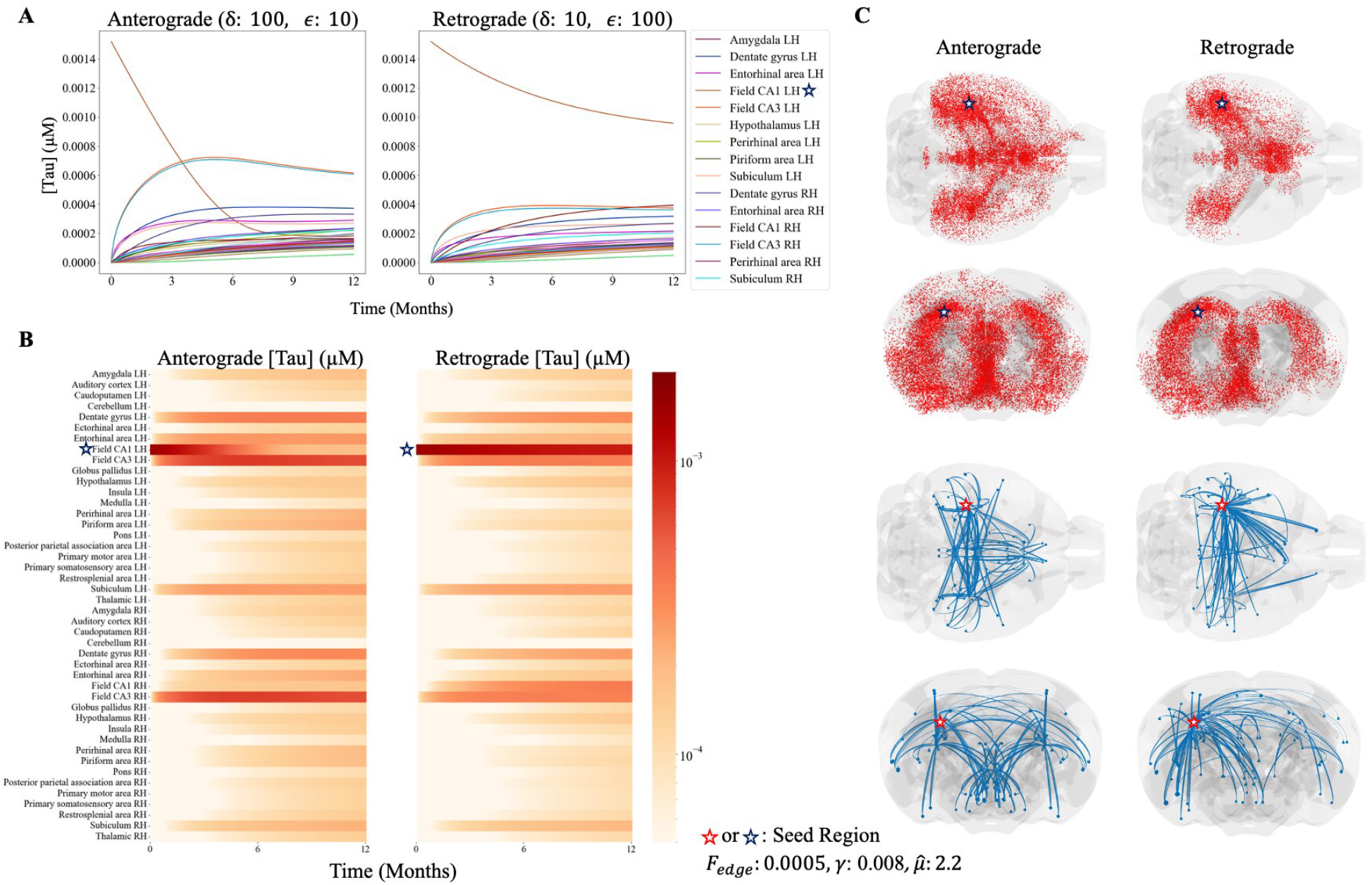
Effect of directed tau transport bias on NTM simulations. **A.** Regional soluble tau concentration over time plots for a 44 region parcellation of the mouse brain (converted the 426 region MCA atlas used in NTM simulations) under anterograde (left; *δ* = 100, *ϵ* = 10) and retrograde (right; *δ* = 10, *ϵ* = 100) conditions. **B**. Heatmap representation of per-region soluble tau concentration over time shown in **A. C**. Glass mouse brains displaying the spatial distribution of soluble tau in the brain (top two rows) at 6 months post seeding under anterograde (left) and retrograde (right) biased transport regimes. Higher regional density of red dots indicates a higher concentration of tau. Glass brains showing the top 0.5% of interregional tau fluxes are shown with blue arrows (bottom two rows). The seed region (hippocampal subfield CA1, left hemisphere) is indicated by a red or blue star.

*In contrast with γ*, modulating *δ* and *ϵ* parameters has a critical influence on the spatial pattern of tau spread. In the anterograde regime (*δ* = 100, *ϵ* = 10), the tau trajectory of the seed region indicates a much faster spread to other brain regions (Figure 4A,B; left) compared to the retrograde regime (right), where the seed region maintains a soluble tau concentration more than 2.5 times the concentration of the next most tau-dense region throughout the duration of the simulation. The order of most to least tau-concentrated regions shows a stark difference between anterograde and retrograde regimes as well, with hippocampal subfield CA3 in the left and right hemisphere exhibiting the highest tau concentrations after 4 months (where they eclipse the seed region), but is much lower, and eventually eclipsed by hippocampal subfield CA1 in the right hemisphere as well as remaining much lower than the seed region (CA1, left hemisphere), in the retrograde biased transport condition. This indicates that modulating the dynamics of active transport has a strong impact on the staging of tau progression (i.e. its spatial pattern) on the macroscale level. Examining the largest inter-regional tau fluxes at 6 months (panel C; bottom two rows) reveals notably different flux patterns between anterograde and retrograde transport regimes. In the anterograde condition, flux maps appear to be nearly symmetric across left and right hemispheres, while in the retrograde condition there is a notably larger source of flux coming from the seed region (CA1, left hemisphere) and spreading to other regions within the left hemisphere as well to across hemispheres. In the retrograde condition, there are comparably fewer large flux vectors found contained within the right hemisphere.

### 3.3 Varying cellular tau uptake and release rates

Modulating the uptake and release rate of tau between intracellular environment of neurons represented by connectome graph edges and the extracellular environments of regions on graph nodes shows a strong tie to the global spread rate of tau as captured by the NTM. Tau trajectories over time (Figure 5A, B) for four different uptake and release rates 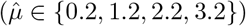 demonstrate that increasing the rate of tau uptake and release increases the global spread rate of tau, increases the rate at which crossing occurs, allowing for faster spread of tau along the structural connectome. It thus appears to have the effect of scaling the time axis of the simulation, where higher values of 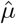 cause faster simulations. By having this effect on the rate of global diffusion, the uptake and release dynamics via 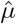 govern when each region’s peak tau value occurs. However, 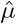 does not appear to impact the ordering of regions in terms of highest to lowest tau concentration. The global tau burden also does not appear to be affected by 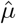, which is again unsurprising.

**Figure 5.**
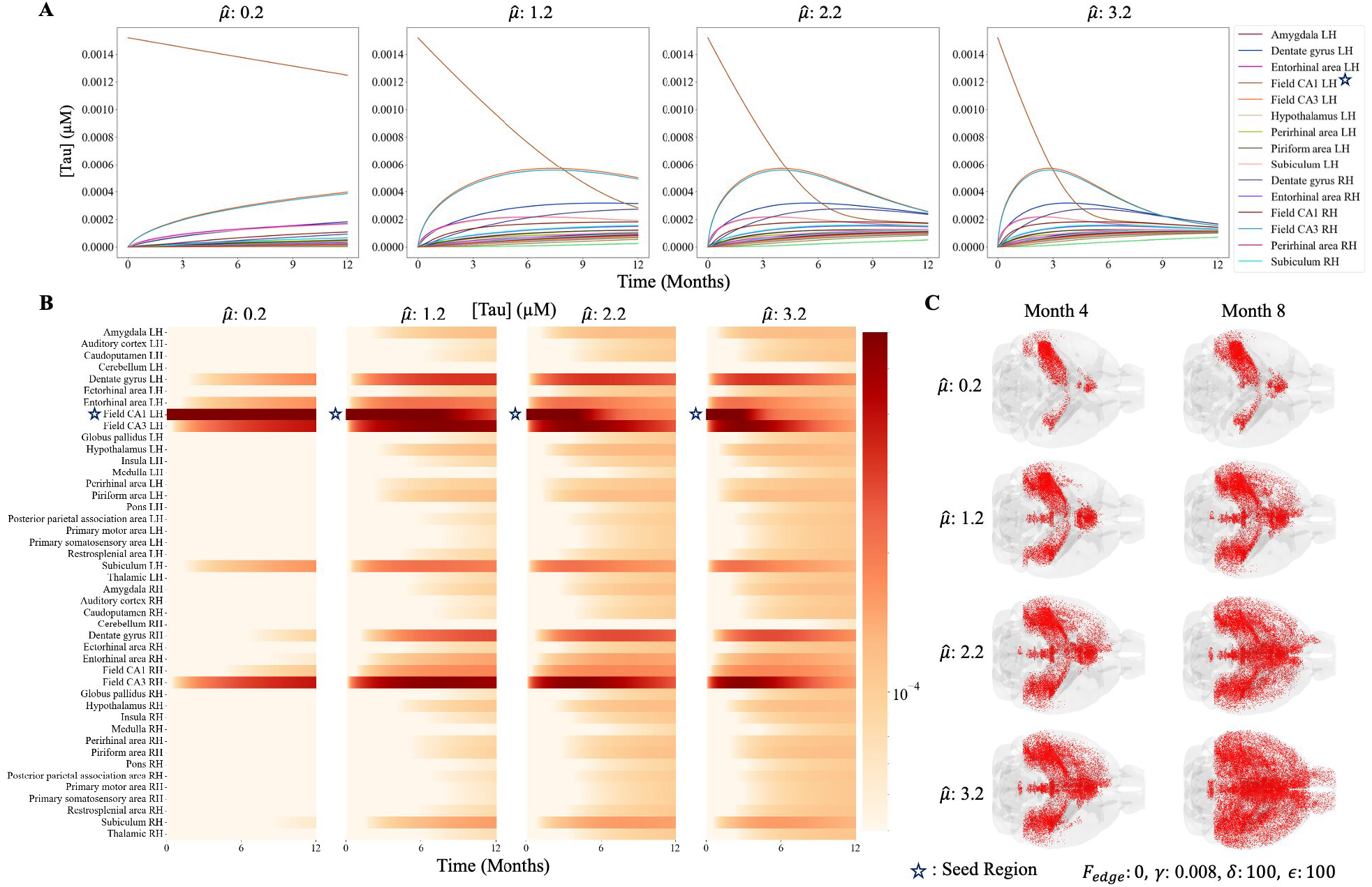
Effect of cellular uptake and release of tau on NTM simulations. **A.** Soluble tau concentration over time plots for a 44 region parcellation of the mouse brain (converted from the 426 region MCA atlas used in NTM simulations) under increasing (left to right; increasing 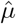) tau uptake and release conditions. **B**. Heatmap representation of per-region soluble tau concentration over time shown in **A. C**. Glass mouse brains showing spatial distribution of soluble tau in the brain at 4 months (left) and 6months (right) post seeding under increasing cellular tau uptake and release conditions (top to bottom). Axial (top) slices are shown. The seed region (CA1, left hemisphere) is denoted with a blue star.

### 3.4 Varying seed-region tau production rates

Increasing the production rate of tau at seed regions is tied most strongly to the global tau burden, as demonstrated by Figure 6. Seed production rate of tau is modulated by the *F*_*edge*_ parameter in the NTM. When this parameter increases, all regions of the brain show an increase in tau concentration in tau trajectories over tau (Figure 6A) and in glass mouse brains showing regional distributions of tau concentration at select time points (Figure 6B). This is expected, as the production of tau is directly responsible for increasing the amount of tau present in seed regions spontaneously over time, which then spread to the rest of the brain.

**Figure 6.**
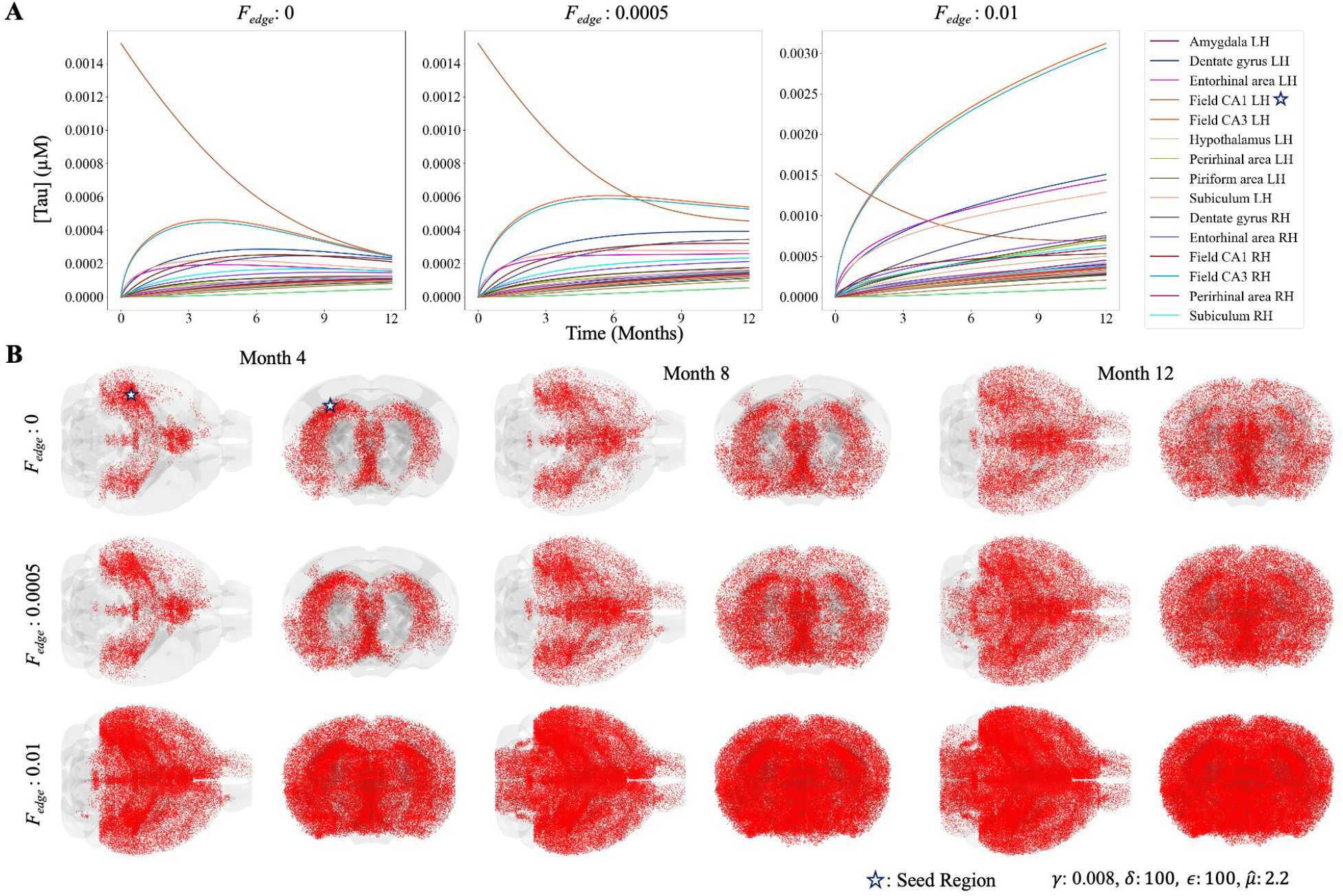
Effect of tau production on NTM simulations. **A.** Soluble tau concentration over time plots for a 44 region parcellation of the mouse brain (converted from the 426 region MCA atlas used in NTM simulations) under increasing (left to right; increasing F) tau production conditions. **B**. Glass mouse brains showing spatial distribution of soluble tau in the brain at 4 month intervals post seeding under increasing tau production conditions (top to bottom). Axial (top) slices are shown. The seed region (left CA1) is denoted with a blue star.

The staging of tau does not appear to be affected except at extreme high values of production, as the ordering of regions in terms of tau concentration appears to remain unchanged when modulating *F*_*edge*_ at lower to intermediate values. The seed region is the one exception, which undergoes a more gradual increase in tau concentration relative to other brain regions as *F*_*edge*_ increases. An explanation for this is that *F*_*edge*_ modulates tau concentration in the intracellular environment of connectome graph edges. This means that intracellular tau production as governed by the NTM occurs in cells connected to both seed regions and all connected brain regions.

### 3.5 Comparison to empirical tau progression

In order to illustrate that NTM simulations of tau progression demonstrate biological accuracy we display empirically obtained spatial distributions of tau from mouse studies (Figure 7) and compare these to simulations generated by the quasi-static NTM. In these studies, mice are injected with tau at a known seed location in the brain, and at specific later time intervals have their brain’s probed from tau abundance in different regions. We display glass mouse brains of empirically derived spatial tau distributions obtained by Iba *et al*. [21] (Figure 7A), referred to here as *IbaHippInj*, and Kaufman *et al*. (Figure 7B), referred to here as *DS6*, collected at three time points after seeding. These empirical tau distributions display a stark resemblance to NTM simulated tau propagation patterns and demonstrate important characteristics of tau spread patterns captured by the NTM. For instance, the *IbaHippInj* tau progression dataset (Figure 7A) resembles NTM simulations displayed in Figures2B (right), 4C, 5C, and 6B (where *F*_*edge*_ ≤0.0005), where tau concentration appears most heavily concentrated in the dorsal and ventral hippocampus but show notable increase in concentration within other regions over time. The hemispheric differences favoring the side containing the seed site, which are most prevalent early in empirical data, (Figure 7A,B; Left Column) are also captured by NTM simulations, as seen most visibly in Figure 5C and to a lesser degree in 6B. Empirical tau progression patterns in the *DS6* dataset (Figure 7B) display these characteristics as well, but also demonstrate a notable increase in overall tau burden over time, which is captured by the NTM’s newly introduced production dynamics (6B).

**Figure 7.**
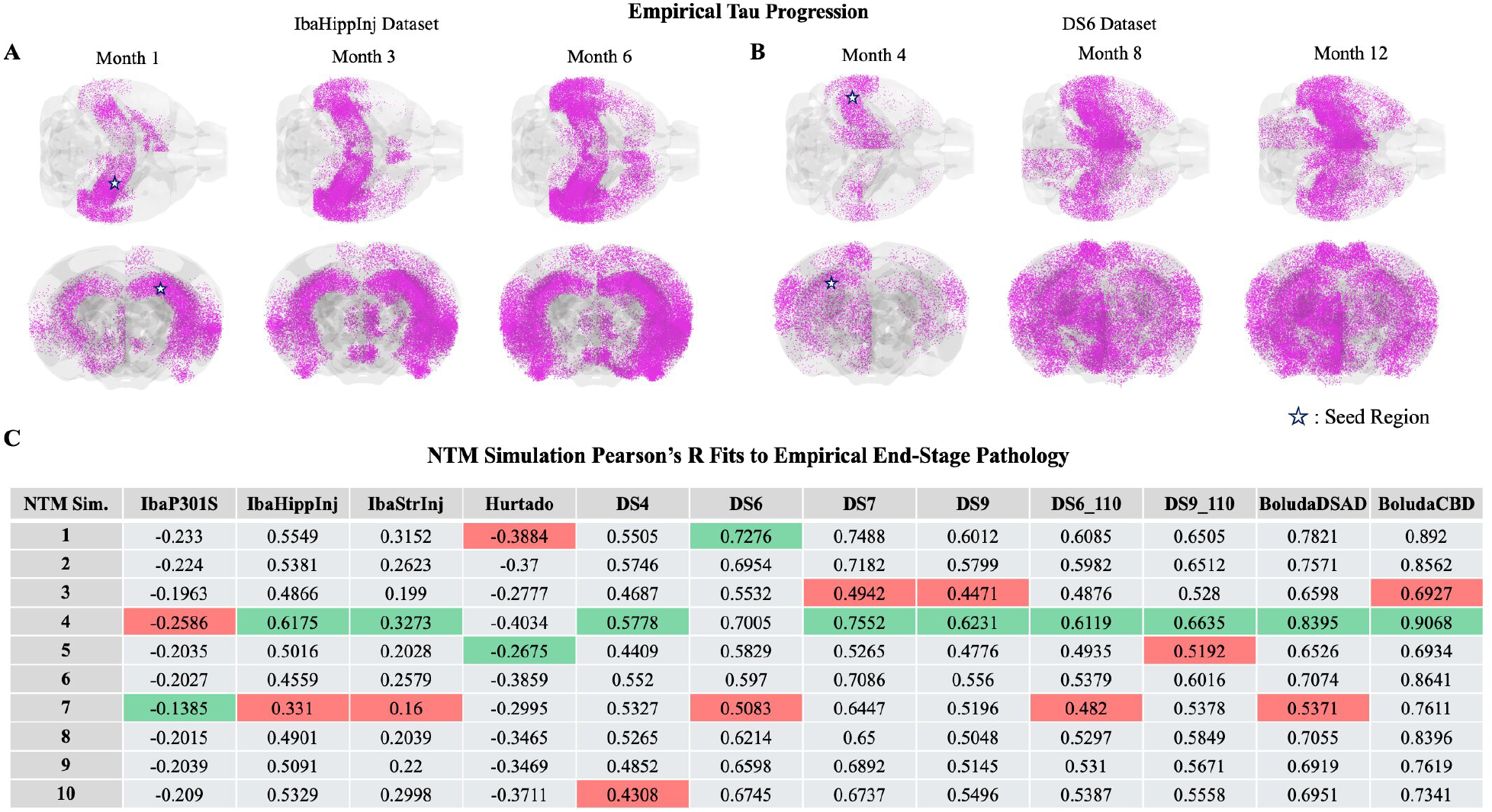
Empirical progression of tau in mouse models and NTM fits. Glass mouse brains showing spatial distributions of tau obtained from empirical mouse studies at sampled intervals. **A.** A mouse study of tau pathology spread obtained from Iba *et al*. [21] referred to here as *IbaHippInj* where tau is seeded in Dentate Gyrus, Right Hemisphere. **B**. A mouse study of tau pathology spread obtained from Kaufman *et al*. [26] referred to here as *DS6* where tau is seeded in left CA1. Increased density of purple dots indicates increased detection of pathological tau. The seed region is indicated with a blue star. (**C**) Pearson’s R correlations between NTM predicted regional patters of tau pathology from the 10 simulations shown in this work compared to 12 empirical mouse datasets at the empirical end stage time point. The best correlation for each empirical dataset is shown in highlighted in green, and the worst performing simulation is highlighted in red. Parameter values and seeding conditions for each NTM simulation is specified in Supplementary Table S2.

Fitting the NTM to these datasets to find ideal parameters minimizing the difference between simulated NTM outputs and empirically observed data is currently intractable due to computational limitations. Each NTM simulation on the 426-region mouse connectome takes on the order of 10 hours to compute despite utilizing parallelization optimizations and high performance compute resources. The slow simulation speed comes from the complexity of solving the underlying system of differential equations defining the quasi-static edge problem multiple times at each time step over a large number of connectome network edges in order to calculate the network fluxes, in addition to integrate the differential equation defining the NTM on the network nodes over a fine grid of discretized time points. Approaches to determine ideal model parameters to fit to data generally take thousands, or orders of magnitude more, model forward simulations, which make parameter inference intractable with the current framework. However, we demonstrate the NTM model’s ability to capture empirical data by comparing the simulations used for biophysical exploration in this study to empirically observed tau progression patterns obtained from mouse studies. The 10 simulations used to demonstrate the effect of perturbing kinetic tau parameters are defined in Supplementary Table S2. The Pearson’s R correlation between NTM predicted spatial patterns of tau and empirical patterns of tau at the end-stage of 12 empirical mouse datasets of tau progression are shown in Figure 7C. These results show that the NTM’s ability to capture end stage spatial tau patterns reaches as high as Pearson’s R of 0.91 even without advanced approaches to identify ideal model parameters. This analysis also demonstrates variability between the NTM simulation fits to empirical data across empirical studies and model parameters. These comparisons serve to demonstrate that the rich diversity of tau propagation patterns captured by NTM simulations match biologically observed patterns of tau progression.

The end-stage tau correlation with empirical data reported here for NTM simulations achieve r-values generally slightly lower than those previously reported by NexIS:dir, an alternate, less biophysically accurate diffusion based models of directionally biased tau spread on the structural connectome [50]. Across the 9 empirical datasets used here with seeding in or near Field CA1 of the left hemisphere of the mouse brain, NTM demonstrated an r-value that was lower by 0.04 than reported by NexIS:dir on average. Notably, the r-value the NTM achieves is nearly equal or notably higher in 3 out of 9 empirical datasets used here (namely *DS7, BoludaDSAD*, and *BoludaCBD*), as shown in Figure 8B. Note, however, that the NTM r-values are obtained from a sparse sampling of 10 simulations in different biophysical regimes of independent microscopic kinetics modulating tau spread, while the models against which NTM is benchmarked use sophisticated parameter inference techniques to identify ideal model parameters to fit to data. We also compare NTM fit metrics against the simpler NDM [37, 38] and NexIS:global [4] models at all empirical time points (as opposed to end-stage) in three datasets, as shown in Supplementary Table S3. For DS6 and DS9 datasets, the NTM obtains a fit metric intermediate to those the NDM and NexIS:global models report [4], and substantially improved on both for the DS4 dataset, even without fitting ideal model parameters for the NTM.

**Figure 8.**
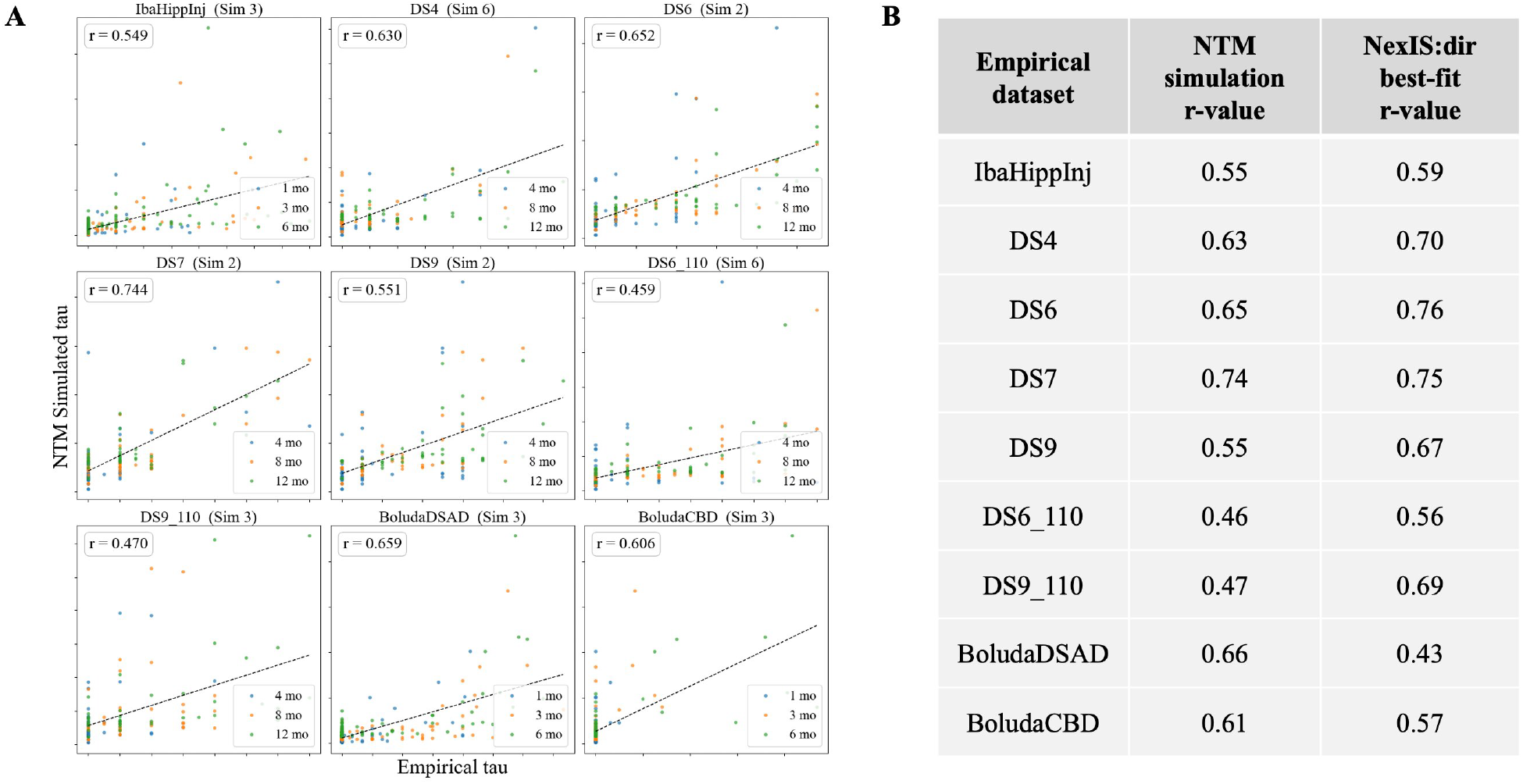
Benchmarking the NTM against alternative models. **A.** Scatterplots showcasing NTM prediction accuracy across brain regions and all sampled time points (as opposed to end-stage spatial distribution of tau pathology; see Figure 7) against 9 empirical datasets of tau progression in P301S mouse studies with tau injectate seeding in or near Hippocampal Field CA1. Note that the *IbaP301S, IbaStrInj*, and *Hurtado* empirical datasets are excluded here because the tau seed region used in the empirical studies these datasets originate from are different. For details see Supplementary Table S1. **B**. Comparison of Pearson’s r-values between model-predicted and empirical tau abundance. The NTM r-values are chosen as the highest across the 10 simulations used for analysis of biophysical parameters without fitting ideal NTM model parameters to each empirical dataset. The NexIS:dir model r-values are computed after fitting the model parameters to the empirical dataset as reported in [50]. NTM simulation numbers correspond to those in Figure 7; See Supplementary Table S2 for details.

## 4 Discussion

The modeling of tau protein spread via the brain’s structural connectome has gained credence in recent years and has proven useful for testing theories of pathological progression of tau on the brain’s structural connectome as seen in Alzheimer’s and other tauopathies without the need for costly and prolonged experimental studies. Network Diffusion based models have demonstrated success in capturing empirically observed pathology progression and disease staging in mouse [31, 32, 19] and human studies [37, 48, 28, 17, 39, 51]. These models have been used to uncover candidate biomarkers associated with tau spread in the brain [17, 24, 42, 55, 54, 8] and for AD staging and subtyping [52]. However, recent investigation has emphasized the important role of microscopic biophysical mechanisms in neurons including active axonal transport of tau by molecular motors acting on the microtubular cellular scaffolding in anterograde and retrograde directions relative to neural polarity [43], interactions of tau hyperphosphorylation, motor proteins and microtubules, as well as aggregation and fragmentation dynamics, tau uptake and release between intracellular and extracellular environments, and trans-synaptic propagation of tau [15, 40, 45, 2, 3, 29].

The present model was devised to bridge the gap between macroscale network-level spatiotemporal progression of tau and the microscale biological mechanisms from which these patterns arise, a facet rarely addressed by earlier models. Our work follows and expands upon the originally proposed Network Transport Model (NTM) [46], which developed a network-wide implementation of aggregation and transport dynamics in a single neuron that was proposed by Torok, et al. [49] to model tau dynamics along brain structural connectome edges. Here we simulate tau spread on the whole mouse brain, instead of prior studies [46] where only smaller subnetworks of the brain were considered for computational tractability. Generating whole brain simulations of tau spread provides not only a more complete image of how tau propagates, but also provides more specific insights, such as how relative spread patterns within versus between hemispheres compare under different biophysical conditions, which could not be drawn from previous localized simulations.

Although the prior NTM [46] provided a preliminary model linking microscale cellular dynamics of tau to macroscale spatiotemporal patterns of tau spread, it did not capture three key dynamics important in mediating tau spread in the brain: 1) the exchange of tau between intracellular and extracellular environments mediated by cellular uptake and release; 2) the continuous production of toxic tau; and 3) a biophysically interpretable mechanism that can control the relationship between slow and fast timescales - a key puzzle in the field [43, 49, 7, 46, 36]. Introducing these biophysical mechanisms into the present NTM model incorporates previously unseen dynamics in simulated outputs of tau progression. By compartmentalizing intracellular and extracellular environments the present model allows for processes such as tau production and interactions with glial cells that are known to occur preferentially or exclusively inside or outside of neuronal cells [18, 1, 9]. To bridge another gap, between biophysical complexity and computational feasibility, we derived a quasi-static approximation that captures the steady-state edge dynamics while allowing for long-term network evolution, as previously employed [46, 6]. We took advantage of the separation of time scales (i.e. a “fast” time scale for edge dynamics and a “slow” time scale for node level dynamics) to obtain a quasi-static NTM model to simulate tau spread in the brain in a computationally feasible manner.

This study focused on the theoretical validation of a biophysical framework. Numerical simulations were run using the mouse mesoscale connectome obtained from viral tracing methods by the Allen Institute [34]. The proposed NTM with uptake, release, and production dynamics exhibits a diversity in tau propagation patterns that previous graph diffusion or earlier NTM models are unable to capture. Full parameter inference and fitting to empirical data are important goals for future studies but are outside current scope; see Limitations. Instead, by demonstrating that the model generates rich empirically plausible macroscale patterns from microscale principles—independent of empirical fitting—and the model’s ability to capture end stage pathology from empirical studies of tau propagation in mice, we establish the model’s potential predictive power and its utility as a generative tool for the broader neuroscience community. Below we discuss specific explorations of biophysical mechanisms using parameter sensitivity analysis, and their biological implications.

### Mechanistic explorations and insights

#### Aggregation rate

The key role of microscopic tau aggregation rate *γ* was demonstrated in Figure 2 and Figure 3. The microscopic kinetic aggregation rate was found to mediate the balance between the abundance of toxic tau fibrils and the global spread of tau in the brain, two different attributes contributing to the toxicity of tauopathic diseases. Higher aggregation rates increases the amount of polymeric tau filaments, but increased the timescale of network-wide progression as well as overall burden of soluble tau in the brain. Mechanistically, this was due to the enhanced and speedy sequestration of tau into non-spreading insoluble species within neurons. Thus, one possible hypothesis for a therapeutic strategy indicated by this outcome is to target speedy sequestration of intracellular tau rather than indiscriminate reduction of tau aggregation kinetics. Interestingly, *γ* seems to also influence the eventual spatial pattern of spread, despite being a global rate constant.

#### Directional transport processes

Next we assessed the role of directional transport velocity via the interaction terms *δ* and *ϵ*, which, respectively, enhance and attenuate the anterograde velocity along axons as tau interacts with the motor protein kinesin. Simulations of anterograde- and retrograde-biased transport processes led to strikingly different macroscopic spatiotemporal evolution patterns, i.e. the staging of tau (Figure 4). This is an intriguing finding, suggesting a mechanistic means by which different tau species and experimental conditions may undertake divergent spatial trajectories simply via differential yet global and spatially invariant transport kinetics.

#### Trans-neuronal release and uptake rates

Figure 5 showcases model behavior under different uptake and release rates (*µ*), which appears to have the effect of scaling the time axis of the simulation, where higher values of *µ* cause faster spread. While they control the effective rate of tau spread, the uptake and release dynamics do not appear to impact either the global tau burden or the ordering of regions, i.e. the spatial patterns that eventually arise. Thus the mechanistic role of the uptake and release dynamics is to regulate how fast tau can cross between the intra- and extra-cellular environments, making *µ* the microscopic parameter most akin to the macroscopic diffusion rate in prior simpler NDM models [37]. It is noteworthy that kinetic control over uptake and release rates in the model establishes the time scale of tau trajectories. In this way the uptake and release rates act akin to control valves between connectome graph nodes and edges modulating the spread rate of tau. Prior modeling attempts have sought to address the vastly different time scales of macroscopic and microscopic processes by simply introducing an arbitrary scaling constant between them. In the previous NTM model [46], the tau diffusivity parameters *λ*_1_ and *λ*_2_ in barrier compartments demonstrated a similar effect. However, these parameters were intended as phenomenological approximations rather than mechanistically accurate ones. Here we were able to naturally recapitulate this timescale disparity via the bottlenecks represented by the release and uptake processes, which are biologically grounded and amenable to experimental intervention.

#### Continuous tau production

The addition of tau production to the NTM introduces important characteristics of tau trajectories that the previous NTM was unable to capture because it was mass conserving. Here we employed production terms that are far more biologically realistic, as both mouse and human studies have displayed an increase in total tau in the brain over time.

Taken together, all innovations proposed in this model - continuous tau production, release and uptake processes accompanied by background protein aggregation and fragmentation kinetics - appear to contribute in complementary and unique ways to produce rich spatiotemporal dynamics of tau progression in the brain. The anterograde or retrograde transport bias modulated by *δ* and *ϵ* effectuate a strong influence on the staging (i.e. spatial patterning) of tau progression. Aggregation rate *γ* displayed a comparatively minor role on tau staging but a substantial role on tau burden, strongly affecting the balance between soluble tau oligomers, considered as vectors of tau spread in the brain, and insoluble tau polymeric fibrils, which are hallmarks of neurodegeneration. The exosome-mediated release and uptake rates 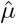 constitute the essential bottleneck of the system, controlling the overall timescale of progression, although not its spatial patterning. Our model may be considered the most mechanistically complete and biologically detailed model to date of endogenous tau progression on the whole brain, not including the role of exogenous factors such as genes and vulnerable celltypes - topics of future explorations.

### 4.1 Comparison to empirical tau progression and alternative connectome-based models

Although the high computational overhead of generating NTM simulations makes parameter inference intractable under the current framework, we validate that the NTM captures empirically observed patterns of tau progression by comparing the simulations used for biophysical analysis with 12 empirical datasets of tau progression from five different P301S mouse studies. For the 8 datasets from studies where tau was injected into Field CA1 in the left brain hemisphere (all but the *Iba* and *Hurtado* labeled datasets), the NTM achieved Pearson’s R values in the range of 0.58-0.91 for predicting the empirical spatial pattern of end-stage tau pathology as shown in Figure 7C. These simulations demonstrate poor accuracy in the *IbaP301S* and *Hurtado* datasets, likely because the seeding of these simulations is substantially different from the seed conditions used for the selected NTM simulations. Despite different seed regions in *IbaHippInj* and *IbaStrInj* datasets, the end stage pathology is similar to NTM-simulated end stage tau (r-values of 0.62 and 0.33, respectively), which can be explained by the fact that the seed regions are nearby and share similar connectivity, so spread dynamics are likely to reach similar end-stage spatial tau patterns. The variance in r-values across NTM simulations for a given empirical datasets shown in Figure 7C also demonstrates the variance in NTM predicted tau patterns as the model’s kinetic parameters vary.

These correlation metrics are generally slightly lower than reported by previously published connectome-based diffusion models of tau spread, but this is to be expected considering that the NTM fit metrics are obtained from a sparse sampling to 10 simulations, while alternative models we benchmark again use sophisticated methods to fit ideal model parameters. Notably, despite the sparse sampling strategy used to obtain an NTM fit metric to real data, it often meets or improves on previous models. On the *DS4* dataset, the NTM predicts tau spread with substantially improved fit metrics compared to the previously published NDM and NexIS models, and obtains an *R*^2^ value intermediate to those obtained by the NDM and NexIS:global on *DS6* and *DS9* datasets (Supplementary Table S2). Comparison of NTM simulation fits to data against the NexIS:dir, a more advanced alternate model of tau spread of the structural connectome, paints a similar story, as shown in Figure 8B, where the NTM achieves r-values generally slightly lower[50], but nearly equivalent or higher in 3 out of 9 datasets we use to benchmark, as seen in Figure 8B. Considering that the NTM achieved these r-values from a sparse sampling of 10 simulations instead being obtained following a more sophisticated approach to fit model parameters to data (as was done with the prior models we benchmark against), we expect these fit metrics to improve with more rigorous parameter inference of the NTM. Despite the sparse sampling approach, it is promising that the NTM simulations achieve similar fit metrics to the prior Nexis model, and even sometimes improved. Overall they demonstrate the NTM’s ability to capture empirical tau progression patterns with similar and sometimes improved robustness compared to prior models of connectome-mediated tau spread, while simultaneously providing stronger biophysical realism.

### Limitations and future work

Although the NTM provides a critical link between global patterns of tau progression and the underlying microscopic cellular mechanisms driving tau spread, there are several important aspects of neurodegeneration related to AD and other tauopathies that the NTM does not capture. For instance, neuroinflammation and neuronal cell death are characteristic of these diseases and have an impact on how tau spreads in the brain [41], but they are not captured by the NTM. Note, cell death generally occurs in later stages of disease, while the NTM aims to model the earlier stages where neuron death is not prevalent as shown by mouse models of disease [14, 21, 26]. Neuroinflammation on the other hand is a complicated process facilitated by microglial cells that has previously been shown to impact both synaptic tau transmission and tau seeding dynamics [9, 5, 13, 23]. Modeling these biological processes is beyond the current scope of the NTM. However, recent work utilizing the Network Diffusion Model (NDM) integrated with gene-associated effects was able to demonstrate that the *Trem2* gene, associated with immune function of microglial cells, has a strong influence on tau spread on the structural connectome network [4]. Future work will incorporate these additional mechanisms into the NTM in a similar manner. Future efforts should also target the role of exogenous factors such as genes and vulnerable celltypes, which may mediate some of the model parameters, that in the current approach are assumed by simplicity to be rate constants, potentially in a spatially variable manner.

In addition to the range of tau propagation patterns demonstrated here there are aspects of the NTM model that we left intentionally unexplored. For instance, all of the simulations in this study enforced equal kinetic uptake and release rates between edges and nodes, but if this constraint were lifted, the NTM would likely exhibit different macroscale dynamics than observed here, given their strong link to the network fluxes. If cellular uptake rates were substantially higher than release rates, then we expect that the intracellular space of graph edges would sequester a greater proportion of soluble tau, which would impact global tau propagation patterns. As another example, in the presented NTM simulations we constrain tau production to occur at the prespecified seed regions, but the NTM can conceivable be run without this constraint, which would likely lead to different tau dynamics. We also decided to set the connectome edge distances to be constant, and let the structural connectivity strength preferentially drive tau transmission along edges. The assumption of constant edge distances is a notable computational simplification to reduce the computational cost and to allow numerical tractability. Each of the constraints we applied were because they were deemed to be biologically reasonable or because they provided a simplification to reduce model dimensionality or clarify the general impact of each of the explored model parameters on global tau propagation patterns. In future work the detailed biological constraints selected for model simulations may change to better suit the investigation at hand. In particular, we will extend our modeling framework to the human brain and we will include realistic variable edge distances, whose effect on the propagation dynamics will be extensively studied.

We refrained from full parameter inference and model fitting to empirical data in this study for two reasons. First, there is a paucity of empirical mouse tauopathy data, which will be resolved over time by accumulating public resources. Second and most critically, while our accelerated quasi-static model represents a sensible reduction in the computational costs compared to [46], it remains computationally expensive and time consuming to compute, due to the computational burden associated with the edge boundary value problem. A single forward simulation on the 426-region MCA mouse brain atlas with fixed parameters takes upward of 10 hours even after the simulation code is optimized and parallelized for high performance cluster computing. Hence full parameter inference on empirical data is currently unfeasible, requiring thousands of model simulations. The same constraint prevents a more thorough sensitivity analysis of the model parameters for a deeper exploration of the impact microscopic kinetic rate parameters have on global tau spread patterns. Future approaches will be centered on computational strategies, such as surrogate modeling, to accelerate simulations and achieve feasible individual-level inference and deeper parameter sensitivity analysis of the NTM.

## Acknowledgments

M.B. and E.C. acknowledge the MIUR Excellence Department Project *MatMod@TOV* awarded to the Department of Mathematics, University of Rome Tor Vergata, CUP E83C23000330006. V.T. has been supported by the grant PRIN 20223R9W7H “Pre-clinical and clinical study on pediatric Traumatic Brain Injury and intranasal Nerve Growth Factor: analysis on cortico-striatal connectivity by network propagation modeling, neuronal tracing and chemogenetics”. N.B. and A.R. have been supported by NIH grants R01AG072753 (A145132), R21AG087921 (A144304), and RF1AG087302 (A144450).

## Code and Data Availability

Code and data for the NTM used in this work is available at https://github.com/Raj-Lab-UCSF/NTM_ru/tree/main.

## 6 Technical Terms

**Structural Connectome** a graph representation of the brain, where nodes represent discrete brain regions (parceled by a specified brain atlas) and edges represent neuronal white matter tract connections connecting region-pairs.

**Tauopathy** a class of neurodegenerative diseases, such as Alzheimer’s disease and frontotemporal dementia, characterized by an abnormal accumulation of misfolded tau-protein in the brain.

**Shooting Method** a method in numerical analysis for solving a boundary value problem by means of a reduction to an initial value problem.

**Quasi-static approximation** a theoretical method to simplify complex, time varying multi-scale systems that assume fast timescales to be instantaneous in slower ones.

