## Supplemental Material for "Biophysically realistic network-level transport model of tau progression with exosome-mediated release and uptake processes"

### FIGURES AND TABLES

| Symbol | Description | Remark |
|---|---|---|
| $x$ | Space | $x \in [0, L]$, where $L$ is the total size of the system in $\mu$m |
| $t$ | Time | $t$, refers to the slow time scale (months) |
| $n_{ij}$ | Soluble intracellular tau | Number of monomeric units per volume within a single set of neurons within edge $e_{ij}$, in $\mu$M |
| $m_{ij}$ | Insoluble intracellular tau | Number of units of aggregates per volume within a single set of neurons within edge $e_{ij}$, in $\mu$M |
| $N_i$ | Soluble regional tau | Number of monomeric units per volume within node $P_i$, in $\mu$M |
| $M_i$ | Insoluble regional tau | Number of units of aggregates per volume within node $P_i$, in $\mu$M |

**Table S1.** Glossary of variables for the Network Transport Model with release and uptake.

| Symbol | Description | Remark |
|---|---|---|
| $D$ | Theoretical diffusivity of $n$ | Estimated to be 12 $\mu m^2$/s* |
| $v_a$ | Native ant. transport velocity of $n$ | Estimated to be 0.7 $\mu$m/s* |
| $v_r$ | Native ret. transport velocity of $n$ | Estimated to be 0.7 $\mu$m/s* |

**Table S2.** List of parameters of the NTM with release and uptake whose values were estimated from previous experimentally derived values Konzack et al. (2007). The parameters have been taken as global, regionally invariant constants.

| Symbol | Description | Remark |
|---|---|---|
| $f$ | Diffusing fraction of $n$ | 0.7 |
| $\beta$ | Fragmentation rate of $m$ | Unimolecular process by which $m \rightarrow n$ ($10^{-6}$) |
| $\gamma_1$ | Aggregation rate, $n + n$ | Bimolecular process by which $n \rightarrow m$ ([0.001,0.008]) |
| $\delta$ | Ant. vel. enhancement factor | Effect modulated by $n$ ([10,100]) |
| $\epsilon$ | Ret. vel. enhancement factor | Effect modulated by $m$ ([10,100]) |
| $\lambda$ | Diffusivity barrier, AIS | Reduces the rate of diffusion within AIS (0.1) |
| $\hat{\mu}_{i,1}$ | release of $n$ at $P_i$ | ([0.2,3.2]) |
| $\hat{\mu}_{j,1}$ | release of $n$ at $P_j$ | ([0.2,3.2]) |
| $\hat{\mu}_{i,2}$ | uptake of $N$ at $P_i$ | ([0.2,3.2]) |
| $\hat{\mu}_{j,2}$ | uptake of $N$ at $P_j$ | ([0.2,3.2]) |
| $F_{edge}$ | production of $n$ on the edge | ([0,0.01]) |
| $F_{node}$ | production of $N$ on the node | ([0,0.01]) |

**Table S3.** List of parameters whose behavior we sought to examine in the present work; the ranges explored are given above. Ant. = anterograde, ret. = retrograde, conc. = concentration, vel. = velocity., AIS = axon initial segment.

| **NTM Sim.** | $\gamma$ | $\delta$ | $\epsilon$ | $\hat{\mu}$ | $F_{\text{edge}}$ | **Figure(s)** |
|---|---|---|---|---|---|---|
| *Sim. 1* | 0.008 | 100 | 100 | 2.2 | 0 | 6 |
| *Sim. 2* | 0.008 | 100 | 100 | 2.2 | 0.0005 | 2, 3, 6 |
| *Sim. 3* | 0.008 | 100 | 100 | 2.2 | 0.01 | 6 |
| *Sim. 4* | 0.001 | 10 | 10 | 2.2 | 0.0005 | 2, 3 |
| *Sim. 5* | 0.008 | 100 | 10 | 2.2 | 0.0005 | 4 |
| *Sim. 6* | 0.008 | 10 | 100 | 2.2 | 0.0005 | 4 |
| *Sim. 7* | 0.008 | 100 | 100 | 0.2 | 0 | 5 |
| *Sim. 8* | 0.008 | 100 | 100 | 1.2 | 0 | 5 |
| *Sim. 9* | 0.008 | 100 | 100 | 2.2 | 0 | 5 |
| *Sim. 10* | 0.008 | 100 | 100 | 3.2 | 0 | 5 |

**Table S4. NTM Simulation Biophysical Conditions.** Biophysical parameter values for the NTM simulations compared to empirical P301S mouse datasets of tau progression. The seed region for all simulations was hippocampal subfield CA1 in the left hemisphere.

| **Empirical Dataset** | **NTM Sim.** | **NDM** | **NexIS:global** |
|---|---|---|---|
| *DS4* | 0.40 (Sim. 6) | 0.06 | 0.17 |
| *DS6* | 0.42 (Sim. 2) | 0.33 | 0.46 |
| *DS9* | 0.30 (Sim. 2) | 0.26 | 0.44 |

**Table S5. Model Comparison.** $R^2$ fit values for the NTM, NDM, and NexIS:global for 3 empirical datasets of tau progression in P301S mice with tau injectate seeded in the CA1 in the left hemisphere. The highest r-value across the 10 NTM simulations described in Table S4 is reported, while the r-values for NexIS:global model are obtained after fitting ideal model parameters to the specific dataset. Fit values for the NDM and NexIS:global are previously reported in Anand et al. (2022).

| **Name** | **Model** | **$ROI_s$** | **Injectate** | **Quantification** | **$n_{ROI}$** |
|---|---|---|---|---|---|
| IbaHippInj | PS19 | $DG_R$ | Synthetic PFFs from 2N4R P301S $\tau$ (T40/PS) and from truncated P301L $\tau$ (K18/PL) | 1,3,6 MPI; MC1 Ab | 102 |
| IbaStrInj | PS19 | $CP_R$, $MOP_R$ | Synthetic PFFs from 2N4R P301S $\tau$ (T40/PS) and from truncated P301L $\tau$ (K18/PL) | 1,3,9 MPI; MC1 Ab | 96 |
| IbaP301S | PS19 | $LC_R$ | Synthetic PFFs from 2N4R P301S $\tau$ (T40/PS) and from truncated P301L $\tau$ (K18/PL) | 1,3,6 MPI; MC1 Ab | 148 |
| Hurtado | PS19/ PDAPP | None | None | 2,4,6,8 mo.; AT8 Ab | 45 |
| DS4 | PS19 | $CA1_L$ | Isolated from AD brain homogenate; prominent nuclear inclusions ("speckles") | 1,2,3 MPI; AT8 Ab | 44 |
| DS6 | PS19 | $CA1_L$ | Isolated from P301S mouse brain homogenate; fibril-like cytoplasmic inclusions ("threads") | 1,2,3 MPI; AT8 Ab | 44 |
| DS7 | PS19 | $CA1_L$ | Recombinant fibrils; prominent nuclear inclusions ("speckles") | 1,2,3 MPI; AT8 Ab | 44 |
| DS9 | PS19 | $CA1_L$ | Recombinant fibrils; prominent nuclear inclusions ("speckles") | 1,2,3 MPI; AT8 Ab | 44 |
| DS6 110 | PS19 | $CA1_L$ | DS6 strain, 1:10 dilution | 1,2,3 MPI; AT8 Ab | 44 |
| DS9 110 | PS19 | $CA1_L$ | DS9 strain, 1:10 dilution | 1,2,3 MPI; AT8 Ab | 44 |
| BoludaDSAD | PS19 | $CA1_L$, $SSP_L$ | DSAD brain homogenate | 1,3,6 MPI; AT8 Ab | 90 |
| BoludaCBD | PS19 | $CA1_L$, $SSP_L$ | CBD brain homogenate | 1,3,6 MPI; AT8 Ab | 58 |

**Table S6. Mouse Tauopathy Datasets.** List of the tauopathy datasets explored here with descriptions of five key experimental features: mouse genetic background, injection site, type of $\tau$ injected, how $\tau$ was quantified, and the number of regions for which $\tau$ pathology was quantified. All $\tau$ pathology was quantified on an ordinal scale. All studies quantified $\tau$ pathology within hemispheres ipsilateral and contralateral to the injection site separately with the exception of Hurtado, which was bilaterally averaged. $ROI_s$ – seeded region; $DG_R$ – right dentate gyrus; $CP_R$ – right caudoputamen; $MOP_R$ – right primary motor cortex; $CA1_L$ – left CA1 region; $SSP_L$ – left primary somatosensory cortex; $LC_R$ – right locus coeruleus; PFF – preformed fibrils; DSAD – Down Syndrome Alzheimer's disease; CBD – corticobasal degeneration; MPI – months post injection; mo. – months of age; Ab – antibody; $n_{ROI}$ – number of regions quantified.
